# A genetically-encoded fluorescent biosensor for visualization of acetyl-CoA in live cells

**DOI:** 10.1101/2023.12.31.573774

**Authors:** Joseph J. Smith, Taylor R. Valentino, Austin H. Ablicki, Riddhidev Banerjee, Adam R. Colligan, Debra M. Eckert, Gabrielle A. Desjardins, Katharine L. Diehl

**Affiliations:** Department of Medicinal Chemistry, University of Utah.; ARC Tangents Analytics; Department of Biochemistry, University of Utah School of Medicine.

## Abstract

Acetyl-coenzyme A is a central metabolite that participates in many cellular pathways. Evidence suggests that acetyl-CoA production and consumption are highly compartmentalized in mammalian cells. Yet methods to measure acetyl-CoA in living cells are lacking. In this work, we engineer an acetyl-CoA biosensor from the bacterial protein PanZ and circularly permuted green fluorescent protein (cpGFP). We biochemically characterize the sensor and demonstrate its selectivity for acetyl-CoA over other CoA species. We then deploy the biosensor in E. coli and HeLa cells to demonstrate its utility in living cells. In E. coli, we show that the biosensor enables detection of rapid changes in acetyl-CoA levels. In human cells, we show that the biosensor enables subcellular detection and reveals the compartmentalization of acetyl-CoA metabolism.

## Introduction

Acetyl-coenzyme A (acetyl-CoA) is a core metabolite that serves central metabolic, catabolic, anabolic, and signaling functions.(1, 2) It is perhaps best known as the intermediate that links glycolysis to the tricarboxylic acid (TCA) cycle. Acetyl-CoA in the cytoplasm is used as the building block for the synthesis of fatty acids and cholesterol, and nucleocytoplasmic acetyl-CoA is the acetyl donor in protein acetylation. In the mitochondria, pyruvate dehydrogenase complex (PDC) converts pyruvate to acetyl-CoA. Acetyl-CoA is also produced from the catabolism of fatty acids in the mitochondria and peroxisomes(3) and of branched chain amino acids (BCAAs) in the mitochondria.(4) In the nucleocytoplasm, ATP citrate lyase (ACLY) cleaves citrate to acetyl-CoA and oxaloacetate, while acyl-CoA synthetase short-chain family member 2 (ACSS2) condenses acetate and coenzyme A to generate acetyl-CoA.(5) PDC can also be present outside the mitochondria and contribute to the nucleocytoplasmic pool of acetyl-CoA.(6–8) Acetyl-CoA produced in the mitochondria (or peroxisome) utilizes the carnitine shuttle via carnitine acetyltransferase (CrAT) to translocate into the nucleocytoplasm.(3, 9, 10) Once the acetyl-carnitine is in the nucleus, carnitine octanoyl transferase (CrOT)(10) converts it back to acetyl-CoA. Acetyl-CoA metabolism is a potential therapeutic target in diseases including cancer,(11–13) metabolic disease,(12, 14, 15) and neurodegenerative disease.(16–18)

Moreover, acetyl-CoA metabolism in bacteria and yeast is of interest since it is the precursor to many valuable chemicals such as isoprenoids, polyketides, and 3-hydroxypropionate.(19–21) Microbial factories offer a sustainable alternative to fossil fuel-derived chemical production.(22–24) Efforts in the field of microbial engineering have focused on maximizing acetyl-CoA in bacteria and yeast to increase the production efficiency of valuable downstream molecules.(19–21) Genetic reporters for malonyl-CoA(25–29) have been utilized in these efforts,(26, 30–33) highlighting the utility of biosensors for enabling metabolic engineering.

Current methods to quantify acetyl-CoA from cells include enzyme-coupled assays (e.g., PicoProbeTM Acetyl-CoA assay) and mass spectrometry.(34–37) Due to the relatively low abundance and poor stability of acetyl-CoA, indirect methods of acetyl-CoA detection are often employed,(38) such as by the analysis of isotopically labeled fatty acids(39, 40) or protein acetylation.(1, 34, 37, 41) Recent advances in quantitating acetyl-CoA and other CoA species from cells by stable isotope labeling of essential nutrients in cell culture (SILEC) and mass spectrometry have achieved subcellular resolution via fractionation of cells and enabled the investigation of acetyl-CoA compartmentation.(34, 35) Nevertheless, these methods, whether direct or indirect, are inherently destructive toward cells or tissues, and subcellular information is dependent on the fidelity of the fractionation technique employed.

Fluorescent biosensors represent a complementary approach for metabolite detection. These sensors are most commonly constructed from a naturally occurring metabolite binding protein and one or more fluorescent proteins (FPs). Biosensors exist for many cellular metabolites (e.g., for ATP, NAD^+^, glucose, etc.) to enable real time detection in live cells,(42–44) but there are only a few examples of acyl-CoA biosensors. For live cell imaging, there is a semisynthetic, FRET-based biosensor for CoA,(45) a ratiometric biosensor for long-chain fatty acyl-CoA esters (“LACSer”),(46) and split luciferase-based biosensor for malonyl-CoA (“FapR-NLuc”).(47) A BRET-based acetyl-CoA biosensor has been reported which can detect acetyl- and propionyl-CoA from cell lysates, but it was not demonstrated to be suitable for live cell imaging in its current form.(48) In this work, we engineered a genetically-encodable fluorescent biosensor for acetyl-CoA and demonstrated the sensor’s utility in live cell detection of acetyl-CoA in E. coli and human cells.

## Results

### Biosensor engineering

We first selected an acetyl-CoA binding protein that we predicted would be amenable to forming a biosensor. We chose the protein PanZ (also known as PanM), which is an endogenous acetyl-CoA sensor that regulates pantothenate synthesis in *E. coli* and other related enterococcus species.(*49–51*) PanZ is a GNAT (GCN5-related N-acetyltransferase) domain-containing protein but was shown to lack acetyltransferase activity.(*49–51*) PanZ binds acetyl-CoA, and this bound state is competent to bind the zymogen PanD and its activated form, aspartate α-decarboxylase (ADC). A structure of the PanZ/acetyl-CoA/PanD complex (PDB: 5LS7) and other biochemical data(*49–51*) suggested that PanZ is a suitable basis for a fluorescent acetyl-CoA sensor. We tested the binding of purified PanZ to acetyl-CoA and CoA using surface plasmon resonance (SPR) and measured a 7-fold higher affinity for acetyl-CoA versus CoA (**Figure S1**). Based on these data, we concluded that PanZ was a promising starting point for our biosensor.

To engineer a fluorescent biosensor, we screened insertion sites of circularly permuted GFP (cpGFP) in loop regions of PanZ (**Figure 1a**). The variants ranged widely in terms of fluorescence intensity (**Figure S2a and S2b**). Considering the fold-change between the +/- acetyl-CoA conditions (**Figure 1b**) and the overall fluorescence intensity of each variant, we selected the R69-E70 insertion site as the best-performing from the panel. We next undertook a linker length screen based on this R69-E70 variant. The initial R69-E70 construct contained an AS linker on either side of the cpGFP (i.e., Nterm-PanZ(1–69)-AS-cpGFP-AS-PanZ(70-137)-Cterm). The linker combinations shown were tested (**Figure 1C**). The GA linker at both the N-terminus and C-terminus of cpGFP was revealed to perform better than the original AS linker by about 2-fold in terms of responsivity (**Figure S2c**). We termed this version of the biosensor PancACe (pronounced “pancake”; for <u>Pan</u>Z <u>A</u>cetyl-<u>C</u>oA sensor; **Figure 2A, S2d**). To attempt further optimization of PancACe, we undertook a linker composition screen (**Figure S2e**). To narrow down which variants to test biochemically, we performed molecular modeling and docking studies (**Figures S3 and S4**). However, none of the linker compositions we tested performed better than PancACe (i.e., GA). Notably, the EE linker variant is a “turn off” sensor. We also tested variants of PancACe in which cpGFP was replaced with yellow (cpYFP, “banana PancACe”) or blue (cpBFP, “blueberry PancACe”) fluorescent protein using the same GA linkers as PancACe (**Figure S4f**). Neither variant performed as well as PancACe, and so we proceeded with PancACe.

**Fig. 1.**
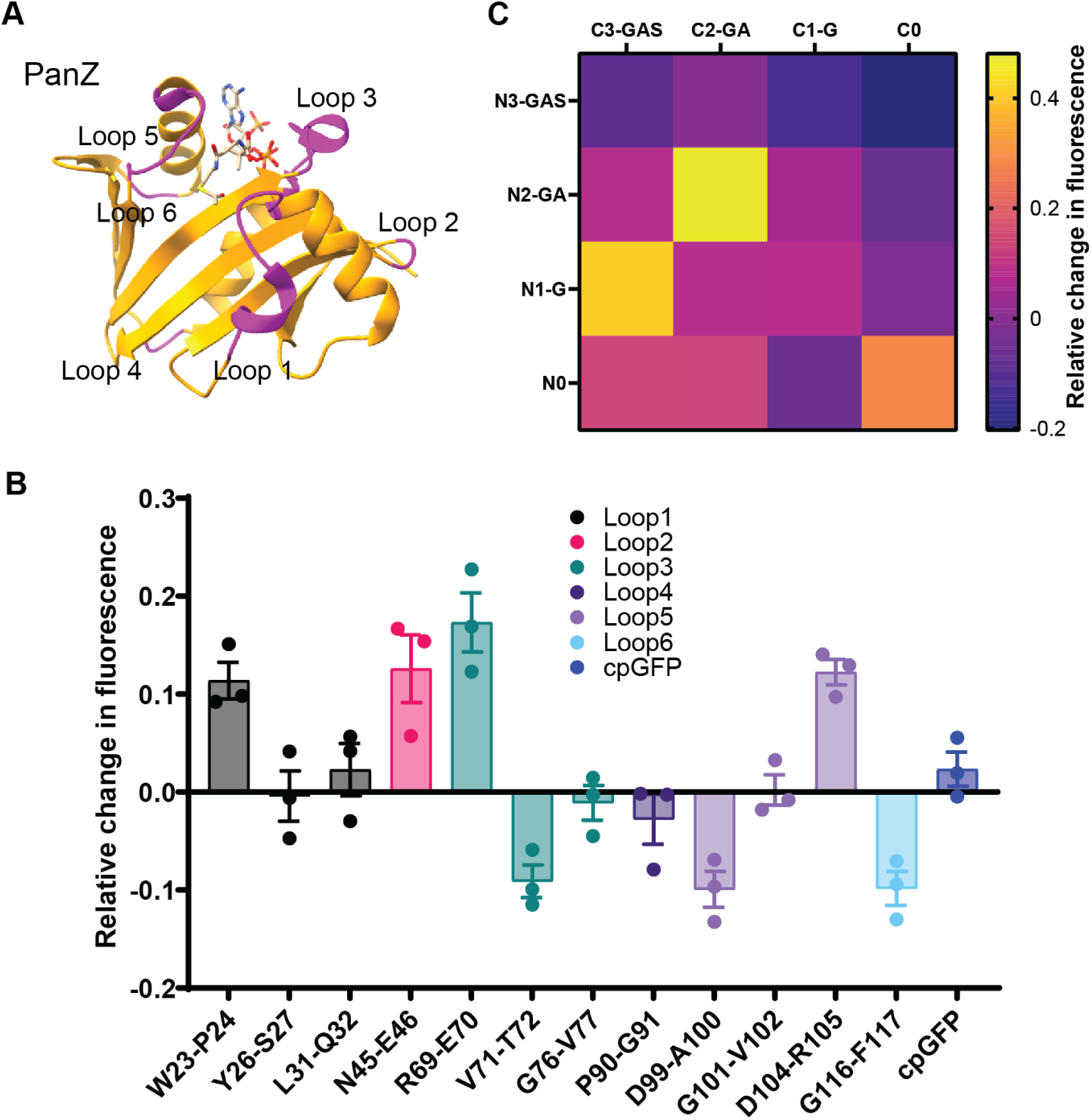
Engineering of an acetyl-CoA biosensor from PanZ. **(A)** Crystal structure of PanZ (PDB: 5LS7) with the loops highlighted (magenta) into which cpGFP was inserted. **(B)** Analysis of the loop insertion mutants. The data are displayed as relative fluorescence in the presence of 1 mM acetyl-CoA compared to in the absence of acetyl-CoA. n = 3. **(C)** Analysis of the linker length variants (PanZ(1-69)-Ntermlinker-cpGFP-Ctermlinker-(70-137)PanZ fusion protein). The data are displayed as relative fluorescence in the presence of 1 mM acetyl-CoA compared to in the absence of acetyl-CoA. n = 1-3. For panels B-C, the data are displayed as the relative fluorescence: (F_1_-F_0_)/F_0_). λ_ex_ = 485, λ_em_ = 528 nm, SEM shown for panel B.

**Fig. 2.**
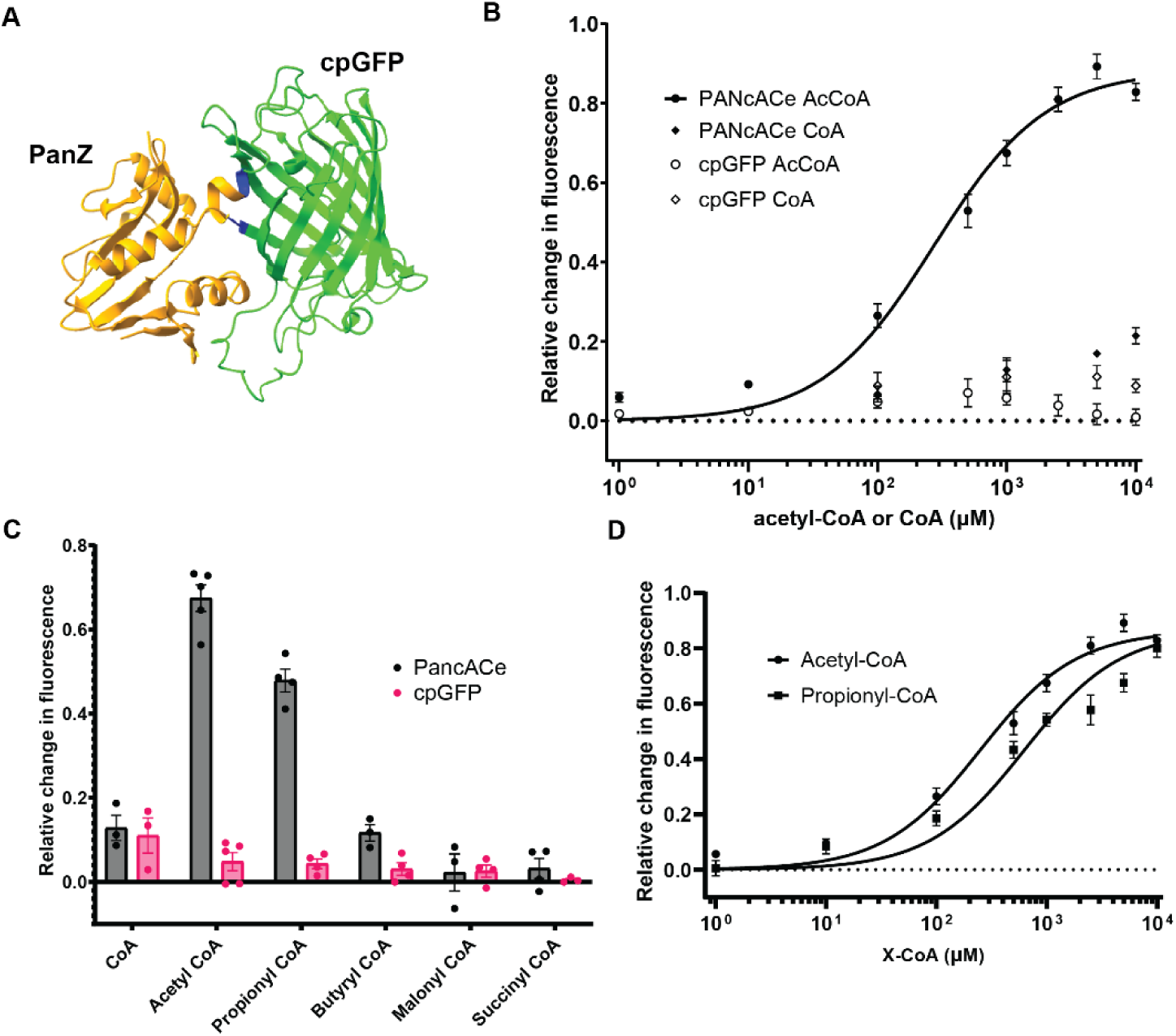
Characterization of the biosensor PancACe. **(A)** Model of PancACe derived from PDB 5LS7 and 3O77. **(B)** Titration of PancACe or cpGFP with acetyl-CoA or CoA. The PancACe + acetyl-CoA data were fitted to a one-site binding model. **(C)** Selectivity testing of PancACe. The data were displayed as relative fluorescence in the presence of 1 mM X-CoA compared to in the absence of X-CoA. **(D)** Titration of PancACe with propionyl-CoA. The data from panel B for PancACe + acetyl-CoA are reproduced on this plot for comparison. The data were fitted to a one-site binding model. For panels B-D, the data were displayed as the relative fluorescence: (F_1_-F_0_)/F_0_). λ_ex_ = 485, λ_em_ = 514 nm, n = 3-5, SEM shown.

### Biochemical characterization of PancACe

We titrated PancACe or cpGFP with acetyl-CoA or CoA (**Figure 2b, S5a**). PancACe displays an increase in fluorescence upon binding to acetyl-CoA and has a maximum response of nearly 2-fold at saturating acetyl-CoA. This response range is comparable to other reported metabolite biosensors derived from a cpFP.(*42*, *52*, *53*) The affinity of PancACe for acetyl-CoA (K_D,app_ = 274 ± 39 µM from fluorescence data, **Figure 2b**) is about 160-fold lower compared to PanZ (K_D_ = 1.7 ± 0.2 µM from SPR data, **Figure S1**). Since the R69-E70 insertion of cpGFP is very close to the acetyl-CoA binding site (**Figure 1a**, loop 3), this effect is not surprising. Indeed, a molecular model of the sensor shows that the cpGFP insertion alters the acetyl-CoA binding site of the PanZ significantly (**Figure S3**). PancACe shows good selectivity for acetyl-CoA over CoA (**Figure 2b**), which is consistent with our SPR binding data for PanZ (**Figure S1**). Even though the PancACe-CoA data cannot be fitted to obtain a K_D,app_ since saturation was not achieved, interpolation between the PancACe-AcCoA and -CoA data indicates that 10 mM CoA is required to elicit the same sensor response as 100 µM acetyl-CoA. We further tested the selectivity of PancACe for short-chain acyl-CoAs (**Figure 2c**). The only other acyl-CoA that was recognized by the sensor was propionyl-CoA. Based on this single concentration experiment, the affinity of PancACe for propionyl-CoA is lower than for acetyl-CoA. A propionyl-CoA titration confirmed this apparent affinity difference (**Figure 2d**). Since saturation was not achieved up to 10 mM propionyl-CoA, we assumed that the same maximal sensor response would be achieved for propionyl-CoA as for acetyl-CoA. Based on this assumption, we fit the data and estimated a K_D,app_ = 626 ± 75 uM for propionyl-CoA, a 2.3-fold lower affinity compared to acetyl-CoA. In most circumstances, there is likely much more acetyl-CoA in cells than propionyl-CoA. However, there are reports of conditions in which propionyl-CoA and acetyl-CoA concentrations are comparable such as in the nucleus.(*34*)

Since fluorescent proteins display pH sensitivity,(*54*, *55*) we measured the fluorescence of PancACe and cpGFP over pH 6.5-8 (**Figure S6**) and observed similar behavior as has been reported previously for an NAD^+^ biosensor derived from cpVenus (cpV).(*42*) Since the pH sensitivity of PancACe and cpGFP are similar, cpGFP can be used in cells as a control for pH as a confounding influence on the PancACe signal. Finally, we sought to determine if we could use the same internal control that was used with the cpV-derived NAD^+^ biosensor.(*42*) That group observed that a second excitation wavelength (λ_ex_ = 405 nm, λ_em_ = 520 nm) of cpV was largely invariant to the presence of NAD^+^ and used this wavelength as an internal control for the amount of sensor present in their cell-based assays. PancACe and cpGFP also have this 405 nm excitation peak that is largely invariant to acetyl-CoA (**Figure S5b**) which we used for normalization in live cell experiments.

### Acetyl-CoA measurements in live E. coli

We next used PancACe to measure acetyl-CoA in live *E. coli*. We expressed either PancACe or cpGFP in *E. coli*, manipulated the nutrient status of the cells, and measured the fluorescence of the cells via flow cytometry or well plate reader. We measured λ_ex_ = 405 nm and 485 nm (both with λ_em_ = 514 nm) to normalize the data (F_488_/F_405_) to account for changes due to pH, protein expression, photobleaching, or other factors that might influence the fluorescence signal but are not indicative of acetyl-CoA.

We reasoned that depriving *E. coli* of glucose and other nutrients would lead to a decrease in acetyl-CoA levels over time compared to cells treated with glucose. Therefore, we incubated cells in phosphate-buffered saline (PBS) for 3 hours and used flow cytometry to measure the fluorescence of the cells. We normalized the signal from each deprived time point to the signal measured in the fed condition, (F_deprived_-F_fed_)/F_fed_ (**Figure S7a**). We used cpGFP-expressing cells that were treated identically to the PancACe-expressing cells as a control. After 2 h of nutrient deprivation, PancACe showed a decrease in its fluorescence response while the cpGFP signal remained invariant over the entire time course (**Figure S7a**). Next, we performed a refeeding experiment in which cells were first incubated in PBS for 3 h and then were refed with 28 mM glucose for 30 min, 1 h, 2 h, or 3 h. In this experiment, we normalized the signal from each refed time point to the signal measured in the deprived condition, (F_refed_-F_deprived_)/F_deprived_ (**Figure S7b**). Within 15 min, PancACe displayed a significant increase in signal, suggesting that the cells recovered their intracellular acetyl-CoA levels rapidly upon refeeding with glucose.

Since the change happened quickly during refeeding, we switched to using a well plate reader to measure the fluorescence *in situ* so that we could take repeated measurements of a cell population in shorter increments. We used injectors to rapidly add a concentrated bolus of the glucose, minimizing the volume change. First, we took cells that had either been pre-starved (no glucose for 3 h) or fed (+ 28 mM glucose for 3 h). Both sets of cells received ammonium as a nitrogen source. After briefly measuring the baseline signal, we added either 28 mM glucose to the deprived cells or buffer without glucose to the fed cells (**Figure 3a, S8**). Only the PancACe-expressing, starved cells that were refed with glucose showed a sharp increase in signal, and this occurred within about 5 min of feeding. Indeed, *E. coli* are able to rapidly exit dormancy, as they are accustomed to experiencing cycles of highly variable growth conditions.(*56*) Interestingly, acetyl-CoA appears to drop slightly from the initial peak before increasing again and reaching a stable level. We do not know what this behavior signifies, but our ability to detect it with PancACe highlights the power of this real-time approach.

**Fig. 3.**
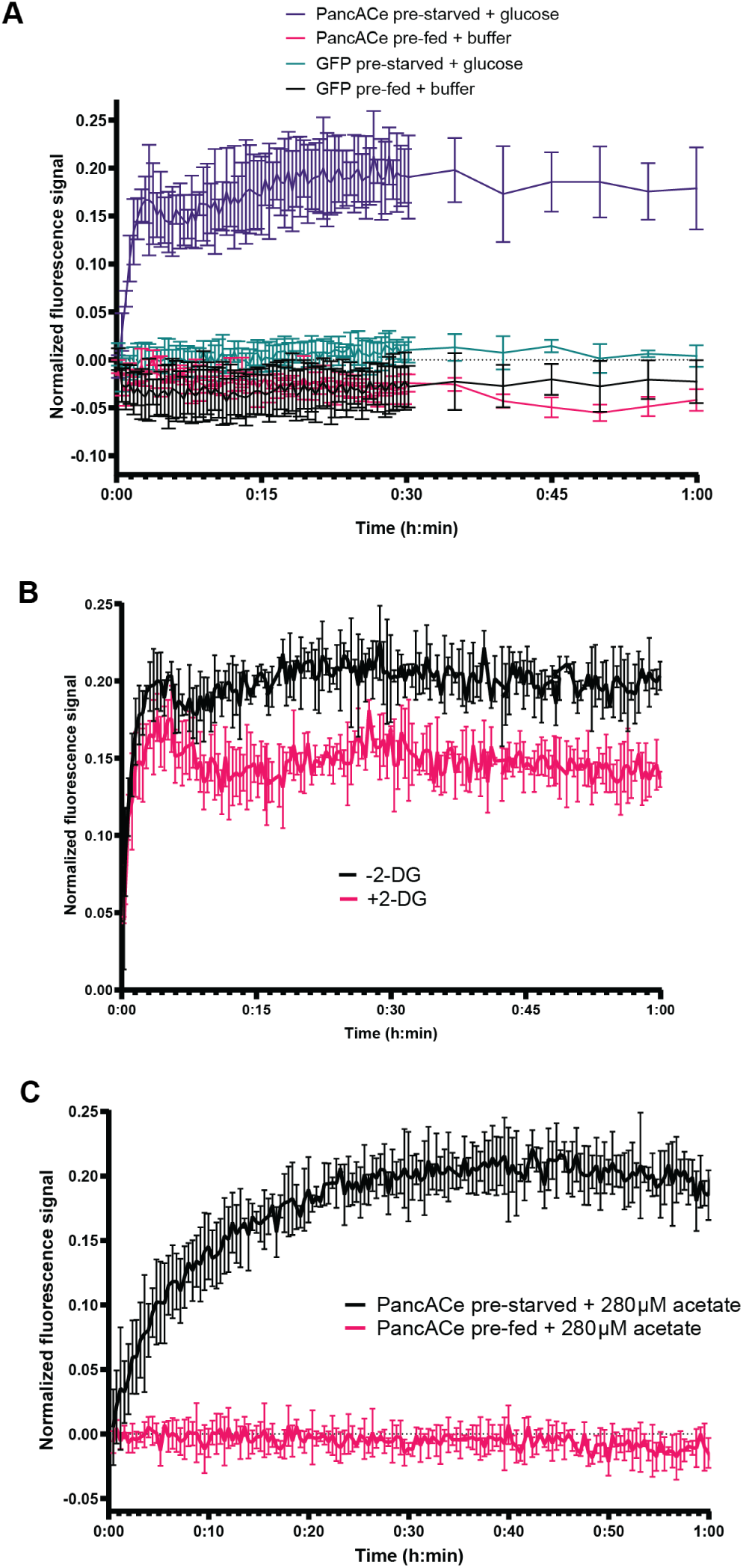
Acetyl-CoA measurements in E. coli by PancACe. **(A)** Normalized fluorescence response of PancACe or cpGFP in *E. coli* to refeeding. The cells were either fed 28 mM glucose for 3 h prior (“pre-fed”) or deprived of glucose for 3 h prior (“pre-starved”). At time 0:00, the cells were given either buffer alone (“+buffer”) or 28 mM glucose (“+glucose”). **(B)** Normalized fluorescence response of PancACe in *E. coli* to refeeding with or without 2-DG. The cells were deprived of glucose for 3 h prior either with or without 2-DG added. At time 0:00, the cells were given 28 mM glucose. **(C)** Normalized fluorescence response of PancACe in *E. coli* to refeeding with acetate. The cells were either fed 28 mM glucose for 3 h prior (“pre-fed”) or deprived of glucose for 3 h prior (“pre-starved”). At time 0:00, the cells were given 280 µM acetate. For panels A-C, the fluorescence was normalized by dividing F_488_/F_405_ then (F_1_-F_0_)/F_0_, where F_0_ is defined by the fluorescence from each population of cells prior to time 0:00. λ_ex_ = 405,485, λ_em_ = 514 nm, n = 3 (technical replicates), SD shown. The additional experimental replicates (n = 3) are shown in **Figures S8-S10**.

Next, we treated cells with 2-deoxyglucose (2-DG), an inhibitor of glycolysis, prior to refeeding. Both groups of cells were deprived of glucose for 3 hours prior to refeeding, and one group also received 28 mM 2-DG during this period. Both groups exhibited an increase in PancACe signal upon refeeding with 28 mM glucose, but the cells treated with 2-DG demonstrated a suppressed increase over the refeeding time period (**Figure 3b, S9**).

Finally, we refed deprived or glucose-fed cells with acetate since acetate can be converted to acetyl-CoA via non-glycolytic pathways.(*19*) Acetate treatment led to a large increase in the deprived condition but no change in the glucose-fed condition (**Figure 3c, S10**). Interestingly, the increase occurred more gradually for acetate refeeding compared to glucose refeeding (**Figure 3a, S8**), supporting that these nutrients are used to make acetyl-CoA via different pathways. We also tested glucose or acetate refeeding on cells that had been fed glucose or acetate for an initial 3-hour period (**Figure S9d**). Only the cells that had initially been fed acetate and were refed with glucose showed a large increase in acetyl-CoA levels according to the sensor. This observation is consistent with glucose being the preferred energy source in these bacteria.(*57*, *58*)

### Acetyl-CoA measurements in HeLa cells

We prepared HeLa cell lines stably expressing PancACe or cpGFP with a cytoplasmic (“cyto”), nuclear (“nuc”), or mitochondrial (“mito”) localization tag (**Figure S11**).(*42*) To determine if the presence of the sensor alters free acetyl-CoA levels in the cells, we used the PicoProbe^TM^ Assay and a histone acetylation immunoblot (**Figure S12**). The PicoProbe^TM^ Assay showed no difference in free acetyl-CoA in the cells expressing cyto-cpGFP versus cyto-PancACe. The blots showed no difference in bulk histone acetylation in the cells expressing nuc-cpGFP versus nuc-PancACe. Together these results indicate that PancACe does not appreciably sequester acetyl-CoA in the cells. To measure acetyl-CoA levels in the cells with the biosensor, we used fluorescence microscopy to quantify the PancACe and cpGFP signals at excitation wavelengths of 405 and 488 nm (**Figure 4a, S11a**) for ratiometric analysis.

**Fig. 4.**
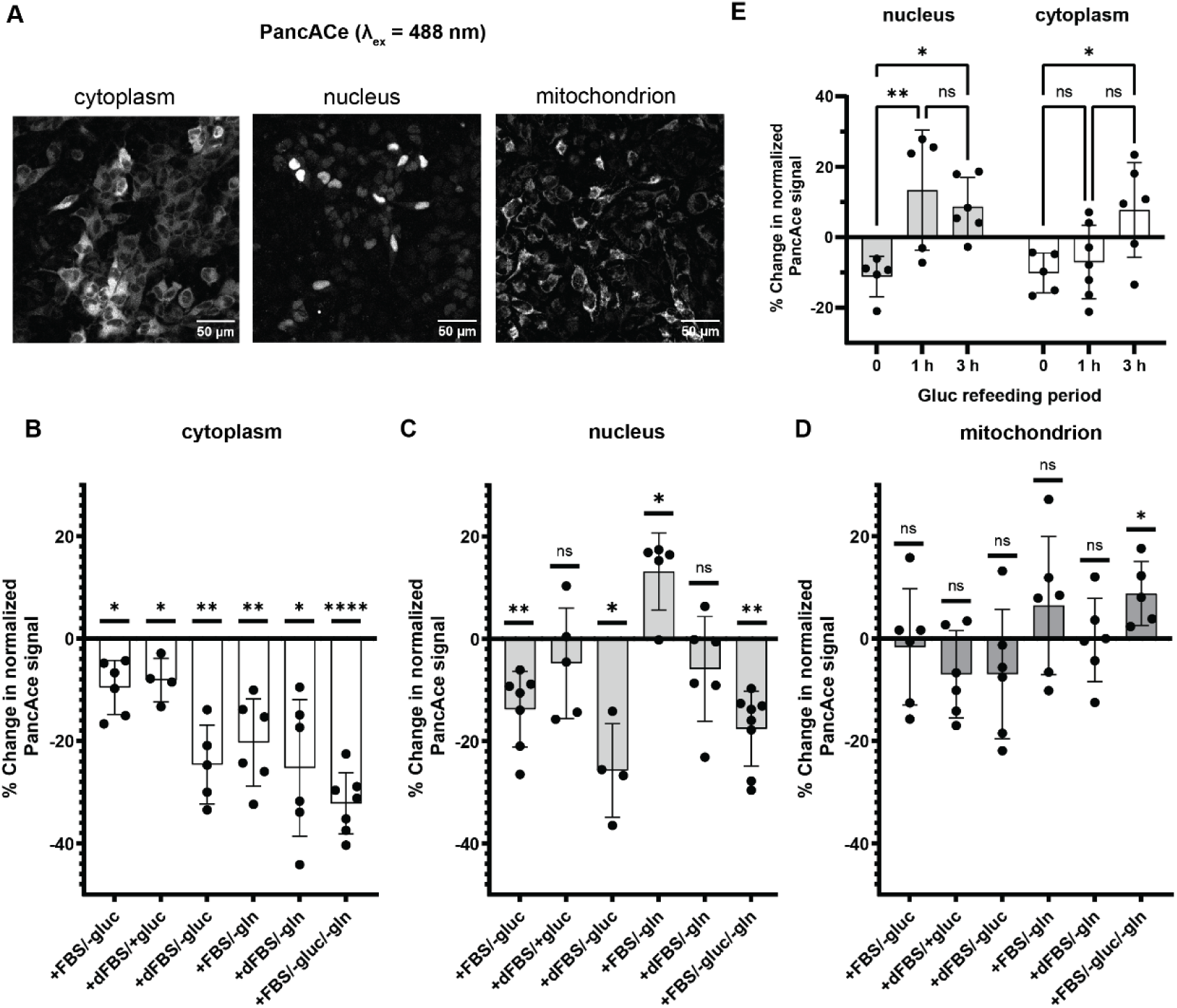
Acetyl-CoA measurements by PancACe in nutrient-deprived HeLas. **(A)** Representative images for each PancACe cell line showing the subcellular localization to each compartment (λ_ex_ = 488). **(B)-(D)** PancACe signal derived from images of HeLa cells expressing nuc-, cyto-, or mito-PancACe that were subjected to the indicated nutrient deprivation for 16 h. The data were collected and normalized as described in the Methods section. The data shown here are normalized to fed cells, n = 4-8 experimental replicates, SD shown. Two-tailed, paired t tests were performed on the data prior to normalization to the fed cells. The p values shown on the plot are for each condition compared to fed cells. e) PancACe signal derived from images of HeLa cells expressing nuc-PancACe or cyto-PancACe that were deprived of glucose for 16 h (time 0) and refed with 4.5 g/L glucose for 1 h or 3 h. The data were collected and normalized as described in the Methods section. The data shown here are normalized to fed cells. n = 4-7 experimental replicates, SD shown. Two-tailed, paired t tests were performed to compare 0, 1, or 3 h to fed cells prior to normalization to fed cells. Two-tailed, unpaired t test were performed to compare 1 and 3 h to 0 h. *p<0.05, **p<0.01, ****p<0.0001. All of the p values are shown in **Tables S1-S2**.

To perturb intracellular acetyl-CoA levels, we deprived the cells of glucose, complete serum (i.e., cells were given dialyzed fetal bovine serum, dFBS), glutamine, or combinations thereof for 16 h. We compared the sensor response to fed cells that were given complete media during the same period. The dFBS was used since normal FBS contains acetate, glucose, and other potential acetyl-CoA precursors(*9*) that dialysis is expected to reduce or eliminate. **Figure 4b-d** display the results for the three cellular compartments tested. Cytoplasmic acetyl-CoA levels were impacted by all of the deprivation conditions. Glucose or serum deprivation alone had a smaller impact than glutamine deprivation or deprivation combinations. The PicoProbe^TM^ assay was used to measure free acetyl-CoA from cells that were glucose or glutamine deprived, and the assay results echoed the decreases reported by cyto-PancACe (**Figure S12b**).

In contrast, nuclear acetyl-CoA levels decreased under glucose deprivation but not serum deprivation alone or glutamine deprivation, although combined serum/glucose deprivation had a larger impact than glucose deprivation alone. Bulk histone acetylation data recapitulated the trends reported by nuc-PancACe (**Figure S13a-d**). In the mitochondria, only the elimination of both glucose and glutamine led to a significant change in acetyl-CoA levels. Interestingly, this condition resulted in an increase in acetyl-CoA compared to fed cells. This behavior could be due to the cells increasing β-oxidation to meet their energy needs in the absence of glucose and glutamine.(*59*)

Next, we tested the recovery of the cells after deprivation. The cells were glucose deprived for 16 h then refed with glucose for 1 h or 3 h (**Figure 4e**). Both cytoplasmic and nuclear acetyl-CoA levels fully recovered within 3 h of refeeding. At 1 h, there was more variability in the nucleus compared to the cytoplasm. Some of the nuclear experimental replicates showed an overcompensation effect at 1 h, suggesting that a surge of acetyl-CoA occurs in the nucleus upon refeeding. This effect was not observed in the cytoplasm at 1 h, indicating a steadier increase in acetyl-CoA upon refeeding in this compartment. When we looked at bulk histone acetylation levels under these same conditions (**Figure S14**), we observed that the levels were beginning to recover after 3 h, which is consistent with a lag between acetyl-CoA level rescue and acetyltransferase activity.

Since PancACe only has a 2.3-fold lower affinity for propionyl-CoA compared to acetyl-CoA, we wanted to evaluate its sensitivity to changes in cellular propionyl-CoA. Previous work showed that branched chain amino acid (BCAA) deprivation lowered nuclear propionyl-CoA levels but not nuclear acetyl-CoA levels.(*34*) In this prior study, the magnitudes of the cytoplasmic reduction in both acetyl- and propionyl-CoA levels were similar under BCAA restriction. We gave cells either complete media or media lacking the BCAAs valine and isoleucine for 24 h prior to imaging (**Figure S15a**). According to PancACe, nuclear acetyl-CoA levels did not change significantly, but cytoplasmic levels dropped to a similar extent as in glutamine or serum/glucose deprivation (**Figure 4b**). Histone acetylation blots showed no decrease in bulk histone acetylation in the BCAA-deprived cells (**Figure S15b,c**). These results are consistent with the prior study’s findings and support that PancACe is reporting primarily on acetyl-CoA and not propionyl-CoA.

Acetyl-CoA is produced and consumed by different pathways in the cell in a compartmentalized fashion (**Figure 5a**). Thus, to measure compartment-specific changes in acetyl-CoA levels using PancACe, we knocked down a panel of proteins (**Figure S16**) that are involved in the production (ACLY, ACSS2, and Pyruvate dehydrogenase E1 component subunit alpha, PDHA), trafficking (CrAT), or consumption (p300 and acetyl-CoA carboxylase, ACC) of acetyl-CoA (**Figure 5b-d**). In the cytoplasm, we observed that the ACLY and ACSS2 knockdowns led to a decrease in acetyl-CoA levels. The CrAT knockdown showed a decreasing trend but was not statistically significant. In the nucleus, we observed that the ACSS2 and PDHA knockdowns decreased acetyl-CoA levels. The ACLY and CrAT knockdowns showed a decreasing trend but were not statistically significant. Knockdown of the acetyltransferase p300 had no statistically significant effect on acetyl-CoA levels in any of the compartments. The cytoplasm and mitochondria showed an accumulation of acetyl-CoA in the ACC knockdown. Knockdown of ACC is expected to reduce fatty acid synthesis and lead to accumulation of acetyl-CoA. We note that the variability in the nuc-PancACe data was too high to determine if the ACC knockdown was statistically significant, although it was trending as an increase. Other than accumulation of acetyl-CoA in the ACC knockdown, mitochondrial acetyl-CoA levels were unaffected by these perturbations. Based on previous cellular fractionation data, mitochondrial acetyl-CoA levels are relatively low compared to nucleocytoplasmic levels.(*34*) One possible explanation for this observation is that mitochondrial acetyl-CoA is converted rapidly into citrate (or acetyl-carnitine for export) rather than accumulating. Therefore, the changes in mitochondrial free acetyl-CoA may be relatively small under the knockdowns. Relatedly, PancACe may not be sensitive enough to detect small changes in an already low level of free acetyl-CoA in this compartment, especially decreases from the baseline.

**Fig. 5.**
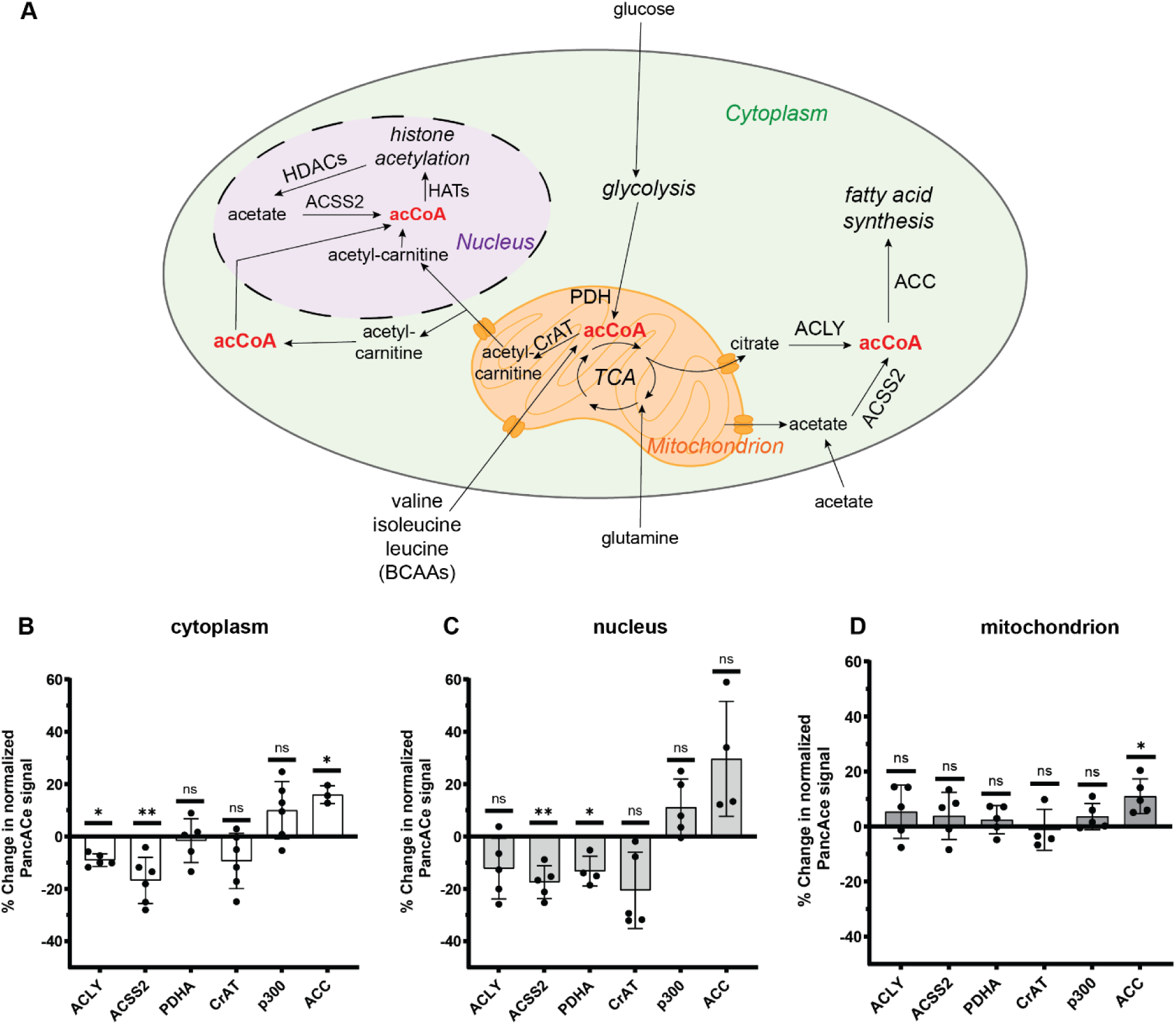
Compartmentalization of acetyl-CoA in human cells. **(A)** Pathways of acetyl-CoA metabolism that we tested with PancACe in the HeLA cells. acCoA = acetyl-CoA, TCA = tricarboxylic acid cycle, ACLY = ATP-citrate lyase, ACSS2 = acetyl-CoA synthetase short chain 2, PDH = pyruvate dehydrogenase, CrAT = carnitine acetyltransferase, ACC = acetyl-CoA carboxylase, HATs = histone acetyltransferases, HDACs = histone deacetylases, BCAAs = branched-chain amino acids. **(B)-(D)** PancACe signal derived from images of HeLa cells expressing nuc-, cyto-, or mito-PancACe that were subjected to the indicated siRNA knockdown for 48 h. The data were collected and normalized as described in the Methods section. The data shown here are normalized to non-targeted siRNA (“NT”) cells, n = 3-6 experimental replicates, SD shown. Two-tailed, paired t tests were performed on the data prior to normalization to the NT cells. The p values shown on the plot are each condition compared to NT cells. *p<0.05, **p<0.01. All of the p values are shown in **Table S3**.

## Discussion

We report herein a fluorescent biosensor for acetyl-CoA that we call “PancACe.” PancACe consists of cpGFP embedded within the acetyl-CoA binding protein, PanZ. As the PanZ domain changes conformation from the unbound to the bound state, the intervening cpGFP is perturbed in its fluorescence emission, leading to an increase in cpGFP fluorescence in the bound state compared to the unbound state (i.e., a “turn on” sensor). Our data show that the insertion of the cpGFP in PancACe leads to significant alteration of the acetyl-CoA binding site compared to PanZ alone. This change results in a large decrease in affinity for acetyl-CoA. Intracellular acetyl-CoA levels in *E. coli* are reported in the range of 20-600 µM,(*60*) while human cells are reported to be in the 1-30 µM range.(*34*, *41*) Based on these figures, PancACe’s affinity for acetyl-CoA is suitable for use in *E. coli*. In human cells, PancACe is likely limited by its sensitivity, particularly in the mitochondria.(*34*) Nevertheless, we were able to detect changes in acetyl-CoA levels across all three compartments in human cells. We further demonstrated that the sensor is responding primarily to acetyl-CoA levels in human cells, suggesting suitable selectivity over propionyl-CoA. Future engineering efforts will focus on producing variants of PancACe with a range of affinities for acetyl-CoA. As is true of other cpFP-based biosensors, the linkers that connect the cpGFP to the PanZ binding domain play a crucial role in determining the responsivity (**Figure 1c**), and perhaps also the affinity, of the sensor. As indicated by our molecular modeling efforts, many challenges still exist with using a computational approach to aid in linker design, highlighting the need in the biosensor field for more refined methods for linker prediction and screening.

A potential application of PancACe is to aid in the metabolic engineering of bacteria or yeast for the production of biofuels or other valuable chemical products. Malonyl-CoA reporters have already been used in this way.(*26*, *30*, *31*, *33*) PancACe’s affinity is well-matched to reported bacterial acetyl-CoA levels.(*60*) We show that the PancACe signal can be monitored in bulk bacterial cultures using a standard well plate reader. The real-time monitoring capability enables detection of rapid changes in acetyl-CoA levels. Alternatively, cells could be sorted by fluorescence to select for subpopulations based on their acetyl-CoA levels as part of a directed evolution workflow.

We further demonstrated that PancACe can be used to report on acetyl-CoA levels in live mammalian cells via fluorescence microscopy. Despite intracellular levels being on the low end of PancACe’s dynamic range, we were able to detect changes in cytoplasmic and nuclear acetyl-CoA levels using ratiometric analysis (ex488/405 and PancACe/cpGFP). We were also able to detect increases in mitochondrial acetyl-CoA. Acetyl-CoA is produced from multiple pathways and precursor molecules, and many of these pathways can compensate for one another.(*9*, *61*) Nevertheless, our sensor was able to detect changes from only partial knockdowns of single proteins involved in acetyl-CoA metabolism. By targeting PancACe to different subcellular compartments, we can observe the compartmentalization of acetyl-CoA metabolism in these cells (**Figure 5a**). Our findings are consistent with the growing body of evidence that the nucleocytoplasm is not a single homogenous pool of acetyl-CoA(*2*, *5*, *6*, *34*, *62–66*) even though acetyl-CoA is small enough to pass freely through the nuclear pore.(*5*) We expect that PancACe and future iterations thereof will be very useful in measuring compartment-specific acetyl-CoA levels in different cell types and potentially in organisms as well.

## Materials and Methods

### Materials

The pET28 vector was a gift from Tom Muir. The plasmid used to subclone the CFP was a gift from Alice Ting.(*67*) The pcDNA3-ER-GCaMP3 plasmid from which the cpGFP gene was subcloned was a gift from Jin Zhang (Addgene plasmid # 64854).(*68*) pLJM1 was a gift from David Sabatini (Addgene plasmid # 19319).(*69*) pMD2.G and psPAX2 were gifts from Didier Trono (Addgene plasmid # 12259 and 12260). Mach1 (Invitrogen) and NEBStable (NEB) cells were used for molecular cloning. Rosetta(DE3) cells (Millipore) were used for protein expression. All acyl-CoAs, including CoA, were purchased from CoAla Biosciences. The 293T cells used for lentiviral production were a gift from Jared Rutter. The HeLa cells used were a gift from Glen Liszczak.

### Molecular Cloning

All PCR was performed with Q5 High-Fidelity DNA Polymerase kit (New England Biolabs) according to manufacturer’s protocols. All primers were ordered from University of Utah DNA/Peptide Core. NEBuilder HiFi DNA Assembly Master Mix (NEB) was used for subcloning. All plasmids were sequence verified by GENEWIZ (Aventa Life Sciences). Amino acid sequences for the genes are listed below (pp 42-26).

#### CFP-PanZ

The *E. coli* PanZ gene was purchased from IDT as a GeneBlock. The PanZ and cyan fluorescent protein (CFP) were subcloned into the pET28 vector including an N-terminal 6x histidine tag followed by a TEV cleavage site (i.e., 6xHis-TEV-CFP-PanZ).

#### cpGFP Insertion Sites Screen

The *E. coli* PanZ gene was subcloned into the pET28 vector including an N-terminal 6x histidine tag followed by a TEV cleavage site (HTSD1.10). The cpGFP gene was supplied by the Jin Zhang lab (Addgene 64854) and subcloned into the pET28 vector using the N-terminal 6x-histidine tag followed by a TEV cleavage site(HTSD1.00). cpGFP was inserted into the PanZ gene between the amino acids shown in **Figure 1b** with an AS linker on the N and C-terminal ends of cpGFP (HTSD6.00-6.13).

#### Linker Length and Composition Screens

cpGFP(HTSD1.00) was inserted into 6x histidine tagged PanZ(HTSD1.10) after amino acid R69 with all 16 possible combinations of N and C-terminal linkers with sequences GAS, GA, G, no linker (HTSD8.00-8.15). Next, the linker composition was varied using Quick Change Mutagenesis: XY = GG, GE, EG, EE QQ, RR, or KK.

#### cpGFP Mutation to Other Color Fluorophores

Quick Change Mutagenesis was performed to cpGFP(HTSD1.00) to generate point mutants cpYFP(HTSD1.01) and cpBFP(HTSD1.02). NEBuilder HiFi DNA Assembly Master Mix(NEB) was then used to insert them into 6x histidine tagged PanZ(HTSD1.10) with GA linkers on the N and C-terminal of the R69-E70 position. This yielded Banana PancACe(HTSD8.05Y) and Blueberry PancACe(HTSD8.05B).

#### Lentiviral vectors

PancACe and cpGFP were subcloned from the pET28 plasmid into the pLJM1 plasmid (Addgene #19319), and the localization tags were added via primers.

### Recombinant Protein Preparation

CFP-PanZ, cpGFP, cpYFP, cpBFP, and all sensor construct plasmids were transformed into Rosetta(DE3) cells for expression. Glycerol stocks of each construct were prepared and stored at -80°C. These glycerol stocks were used to inoculate 10 mL Luer Broth Miller (“LB,” Fisher) media with kanamycin (50 µg/mL). Cultures were shaken at 180 rpm and 37°C overnight then used to inoculate 1 L LB with kanamycin. The 1 L cultures were shaken at 180 rpm and 37°C to OD600 = 0.6. The temperature was then turned to 18°C and flasks were induced with 0.5 mM IPTG for 18 h. Bacteria were pelleted at 4000 x g for 30 min. The pellets were purified or stored at -80°C until purification. Pellet from 1 L of culture was resuspended in lysis buffer (50 mM Tris pH 7.5, 500 mM NaCl, 10 mM imidazole) then sonicated 1 min on/3 min off with duty cycle 50%. Clarified lysates were obtained by pelleting at 17000 x g for 30 min. Clarified lysate from 1 L of culture was incubated with 1 mL of equilibrated Ni-resin (Genscript) and washed with 30 mL wash buffer (50 mM Tris pH 7.5, 50 mM NaCl, 50 mM imidazole). Recombinant proteins were then eluted with 5 x 1 mL aliquots of elution buffer (50 mM Tris pH 7.5, 500 mM NaCl, 500 mM imidazole). Eluents were pooled and buffer exchanged in centrifugal filter units (Millipore; 30kDa MWCO) into storage buffer (50 mM Tris pH 7.5, 150 mM NaCl, 10% glycerol). Proteins were concentrated to ∼100 µM then aliquoted and stored at - 80°C until use.

### Surface Plasmon Resonance

Acetyl-CoA and CoA binding to CFP-PanZ were analyzed via SPR using a MASS-1 instrument (Bruker Daltonics). The CFP tag was used to allow a large portion of immobilization to occur via CFP, thus preserving PanZ’s function. Independent experiments were performed using freshly prepared CFP-PanZ, acetyl-CoA and CoA stocks on two separate days (surfaces 1 & 2). On the same day as the experiment, the CFP-PanZ protein was thawed and buffer exchanged into 1x PBS, pH 7.4 (diluted from 10x PBS, Fisher BP399) using a centrifugal filter unit (Amicon, Millipore Sigma) to remove the Tris storage buffer, which interferes with the capture chemistry. HLC200M sensor surfaces (Xantec Bioanalytics) were activated with EDC/NHS, and then 250 nM CFP-PanZ ligand was captured on the sensor “Spot B” at 10 µL/min at 2020 RU (surface 1) and 2360 RU (surface 2). The surface was then blocked with 1 M ethanolamine. The experimental running buffer was 50 mM Tris pH 7.5, 150 mM NaCl, 0.05% Tween-20, 1 mg/mL BSA for surface 1 replicates and 200 mM Tris pH 7.5, 150 mM NaCl, 0.05% Tween-20, 1 mg/mL BSA for surface 2 replicates. Experiments were performed at room temperature with a flowrate of 30 µL/min. A three-fold dilution series (81 – 0.33 µM) with 1 min association and 2 min dissociation was run in multiple replicates (at least two for each analyte on each surface). Data were double blanked by subtracting the blank in-line control surface (Spot A) and buffer reference injections. Kinetic binding data were globally fit to the Langmuir model for 1:1 binding for each replicate of concentration series with a global R_max_ and fixed global R_min_ (0 RU) using the Sierra Analyser software (version 3.4.1, Bruker Daltonics). The representative replicate data and fit were exported and then plotted using GraphPad Prism version 9.4.1. The average K_D_ from these fits is reported along with the standard error (five replicates for acetyl-CoA and four for CoA).

### Position and Linker Screens

Assays were performed in 96-well plates with a 70 or 100 µL final well volume. All sensors and control proteins were diluted to 10 µM in protein storage buffer (50mM Tris pH 7.5, 150 mM NaCl, 10% glycerol). Acetyl-CoA was dissolved in water at a concentration of 10 mM. A 5x assay buffer was used (1 M Tris pH 7.5, 750 mM NaCl). Wells were set up in triplicate with 60 µL water, 20 µL 5x assay buffer, and 10 µL sensor. Then either 10 µL of water or 10 µL of acetyl-CoA were added before reading. The final concentrations of the assay components were: 1 µM protein, 1 mM acetyl-CoA (or none), 200 mM Tris pH 7.5, 150 mM NaCl. Plates were read at λ_ex_ = 485 nm and λ_em_ = 514 nm in a Biotek Synergy H1 plate reader. Replicates were averaged, and the percent difference between the CoA species and no CoA was calculated: (F_1_-F_0_)/F_0_.

### CoA Titrations and Spectrum Scans

Assays were performed in 384-well plates with a 30 µL final well volume. All sensors and control proteins were diluted to 6 µM in protein storage buffer. Acetyl-CoA, propionyl-CoA or CoA were dissolved in water at a concentration of 60 µM then serially diluted to 30, 15, 6, 3, 0.6 0.06, 0.006 µM. Wells were set up in triplicate with 14 µL water, 6 µL assay buffer, and 5 µL sensor. 5 µL of a CoA stock or water were added before reading. The final concentrations of the assay components were: 1 µM protein, 200 mM Tris pH 7.5, 150 mM NaCl, 0.1 mM DTT, and a variable amount of the CoA species. For point measurements, plates were read at λ_ex_ = 485 nm and λ_em_ = 514 nm in a Biotek Synergy H1 plate reader. Replicates were averaged, and the percent difference between the CoA species and no CoA was calculated ((F_1_-F_0_)/F_0_). For spectra, either the excitation or emission wavelength was fixed, and the other wavelength was varied in 1 nm increments.

### pH Titration

Assays were performed in 384-well plates with a 30 µL final volume. All sensors and control proteins were diluted to 6 µM in protein storage buffer. Different 5x assay buffers were prepared, all 1 M Tris, 750 mM NaCl with pH 6.5, 6.75, 7.0, 7.25, 7.5, 7.75, or 8.0. Wells were set up in triplicate with 15 µL water, 6 µL 5x assay buffer, and 5 µL sensor. The final concentrations of the assay components were: 1 µM protein, 200 mM Tris pH 7.5, and 150 mM NaCl. Plates were read at λ_ex_ = 485 nm and λ_em_ = 514 nm in a Biotek Synergy H1 plate reader.

### Acyl-CoA Screen

Assays were performed in 384-well plates with a 30 µL final volume. All sensors and control proteins were diluted to 6 µM in protein storage buffer. CoA, acetyl-, propionyl-CoA, butyryl-CoA, malonyl-CoA, or succinyl-CoA were dissolved in water at a concentration of 6 mM. Wells were set up in triplicate with 14 µL water, 6 µL 5x assay buffer, and 5 µL sensor. 5 µL of an acyl-CoA or water were added before reading. The final concentrations of the assay components were: 1 µM protein, 1 mM acyl-CoA (or none), 200 mM Tris pH 7.5, and 150 mM NaCl. Plates were read at λ_ex_ = 485 nm and λ_em_ = 514 nm in a Biotek Synergy H1 plate reader. Replicates were averaged, and the percent difference between the CoA species and no CoA was calculated ((F_1_-F_0_)/F_0_).

### Homology Modeling and Energy Minimization

A template was constructed using crystal structures of PanZ (PDB 5ls7) and cpGFP (PDB 3o77). All but one monomer was removed from the PanZ octamer in 5ls7. Next, all but the cpGFP segment of the biosensor was removed from 3o77. Water, acetyl-CoA, and the chromophore fragments were removed from the PanZ and cpGFP molecules. Finally, we made inserts between R69 and E70. The C-terminus at R69 was physically attached to the N-terminus of the linker GA, and the C-terminus of that alanine was attached to the N-terminus of the cpGFP (at the asparagine). Similarly, the C-terminus of the cpGFP was attached to the GA linker, which linked to E70 of PanZ, completing the insertion. Bonds between the respective molecules were created using Dassault Systèmes BIOVIA, Discovery Studio Visualizer, Release [v21.1.0.20298], San Diego: Dassault Systèmes, [2021]. Subsequent models of future constructs were also visualized using the same software. This template was used for homology modelling of the PanZ(1–69)-GA-cpGFP-GA-PanZ(70-137) (PancACe) construct, for which the Robetta server was used.(*70–72*) One hundred models were first generated, from which the top five models were sent back as results (**Figure S4a**). The Ramachandran plots of the five models were then observed and analyzed using PROCHECK™.(*73*) All five models registered favorable residue percentages of 86.2, 89.5, 88.5, 88.8 and 86.8 respectively. A good quality model should have 90% favorable residues or higher. Hence, the second model was deemed the best. The energy minimization was performed using the YASARA energy minimization server, and then analyzed using YASARA View.(*74*) Based on the results of PROCHECK™ and YASARA energies, Model 2 was selected as the de facto model for representing the PancACe construct (**Figure 2a and S4a**).

### Molecular Docking

For observation of all possible interactions of acetyl-CoA with PanZ and PancACe, the SwissdockTM webserver (www.swissdock.ch/docking) was used. First, the PDB file of the protein was submitted to the target selection section of the server. Next, the ligand was selected using Chem3D (or Discover Studio) with all implied hydrogen explicitly shown. Then, the ligand selection was uploaded to the server as a Sybil Mol2 or Charmm file. The webserver operates by docking the ligand to its target in different positions through ligand translation and rotations. The translational variants are designated as clusters, and the rotational variants of the clusters are sub-clusters. Usually, up to 30 clusters are generated for each dock, with up to 15 sub-clusters per cluster. Upon completion, the finished docking file was downloaded and opened using UCSF Chimera, where the protein target and all of the clusters and sub-clusters were displayed. The clusters were selected and opened in Discovery Studio Visualizer to analyze the interactions in 2D.

First, the interactions of the binding site in PanZ with acetyl-CoA were visualized from the PanZ structure (PDB 5ls7; **Figure S4b**). Then Swissdock was used to “back dock” the acetyl-CoA to the same PanZ structure (**Figure S4c-g**). Neglecting the metal interactions, the original crystal structure had the backbone of its interactions centered around a solitary arginine residue (Arg74 in PanZ) which consisted of salt bridges and pi-cation stacks. Also, the salt bridges between the phosphate backbone of acetyl-CoA and surrounding residues were totally utilized. Hence, after the “back dock,” conformations were selected which imitated this binding mode and the acyl-tail orientation of the acetyl-CoA. This led to five clusters which managed to achieve this (**Figure S4c-g**).

The five top clusters were used as the basis to map the changes in the binding site interactions of the PancACe construct with acetyl-CoA (**Figure S4h**). The docking of the PancACe construct with acetyl-CoA immediately revealed that the key anchor residue arginine (Arg 74 in the PanZ numbering) cannot engage with the substrate due to the added bulk of cpGFP pushing the acetyl-CoA out of the binding site, disrupting pi cation stacks and salt bridge interactions. Also, the displacement of the acetyl-CoA from its usual position in the binding site shows underutilization of the salt bridge interactions between the phosphates of the acetyl-CoA and surrounding residues.

Since the chromophore in cpGFP is a fusion of amino acids, the model cannot represent those fragments very well, hence, it has rewritten the sequence in a way that the amino acid fusion is not present. However, since all the interactions are shown in the PanZ fragment, the cpGFP is only there to represent a steric barrier to differentiate from native PanZ-substrate interactions. The steric barrier is significant enough to create more than a hundredfold decrease in binding.

### Homology Modeling and Molecular Docking for Linker Composition

Homology models of PancACe varying the amino acid composition of the linkers (i.e., PanZ-XY-cpGFP-XY-PanZ) were created and used for molecular docking. The theoretical number of linker permutations was too cumbersome (400 variants if both linkers are the same), so the first amino acid was fixed as Gly (X = G), and the second position was varied to all 20 amino acids other than Ala. The SWISS MODEL server was used.(*75–78*) The PancACe model (**Figure** ; XY = GA) was used as a template. All 19 models returned acceptable Ramachandran plots, with favorable residues varying between 90.5% to 91.4%. All models were then energy minimized and analyzed with respect to the XY = GA construct as a reference (-196993.7 kJ/mol YASARA energy, **Table S10**). The top hits were XY = GE or GQ).

Next, the positions of GQ and GE were swapped to EG and QG, along with AG. Linker repeats were also tested including AA, EE, and QQ. The results gave three new top linkers: XY = EG, EE and QQ (**Table S11**). The final top four molecules selected were EE, EG, GE, and QQ. It was noted that the EE construct had a major jump in YASARA energy by 2742 kJ/ mol compared to the parent GA molecule. The change could be primarily attributed to solvation effects but warranted further investigation.

A closer examination of the linkers in the top four molecules showed some glutamate residues actively entering the beta barrel of the cpGFP, especially in the case of the EE construct (**Figure S5a**). This led to an investigation of the mechanism of the chromophore, and how the equilibrium works in favor of the charged/ uncharged chromophore. A careful survey of the mechanistic pathways of chromophore excitation in cpGFP(*79–82*) led to the conclusion that an equilibrium does indeed exist between the green anionic species and the neutral uncharged species, followed by a slow equilibrium between surrounding residues. The neutral and anionic (green) chromophores shuttle back and forth between each other depending on the stability of each species, and the neighboring participation, especially from residues such as E222, which plays a stabilizing role(*80*) Also, one needs to account for the possibility of fluorescence quenching by water, possibly due to hydrogen bonds between anionic chromophores and water being broken and replaced by covalent bonds.(*83*) Similarly, in the structure of the EE construct, two glutamate residues were away from the beta barrel, while two were nested inside it. Hence, we propose that perhaps the two glutamate residues can help in removing some water molecules away from the beta barrel, reducing some of the “quenching load” that the solvent imposes on the cpGFP segment. This could be combined with the possibility that the two glutamate residues inside the beta barrel could contribute to creating a more negatively charged environment inside the beta barrel, forcing the production of more anionic chromophore species *ala* a le Chatelier effect, thereby enhancing the fluorescence signal in the process.

We next decided to also use a traditional docking-based approach to observe whether the linkers can play any active role in improving PancACe binding to acetyl-CoA. First, SWISSDOCK(*84*, *85*) was used to document the reference docking score of acetyl-CoA to the native PanZ. To avoid complications, solvent-free docking was used to examine the interactions of acetyl-CoA with the variants. The PanZ-acetyl-CoA solvent free binding energy was -18.38 kcal/mol. The docking obtained using SWISSDOCK approximated the binding mode from the PanZ crystal structure (**Figure S4b**). Only two binding site residues bound incorrectly to acetyl-CoA compared to the PanZ crystal structure. The approximation was used as a reference for screening the docking of all the linker composition variants. PancACe (XY = GA, **Figure S4h**) was next docked with acetyl-CoA and yielded a binding energy of -14.2 kcal/mol, confirming the biological results of worse K_D_ compared to PanZ.

The docking scores of six linker variants were analyzed, based on top stable models after energy minimization (**Tables S10 and S11**), along with GG as a control. GG was used as it represents the ultimate flexible linker with no steric interference regarding surrounding residues and acetyl-CoA. The binding energies of the top clusters of all linker variants showed that they were either not as good as the XY = GA parent molecule or were slightly better in some cases (**Table S12**). We decided to corroborate the results by biological screening of the linker variants, to prove this. Hence, based on the computational screening, we identified the top four optimal linkers as EE, EG, GE, and QQ. Along with GG (as a control), these four were selected for expression and biological testing for fluorescence.

### Flow Cytometry Deprivation Experiments

All assays were performed with the BD FACSAria III Cell Sorter. All sensors and control proteins were expressed in Rosetta(DE3) cells. PanZ was used as a non-fluorescent control, cpGFP was used as a fluorescent control, and PancACe was the experimental condition. Cultures were grown in LB with kanamycin overnight at 180 rpm, 37°C then used to induce new cultures. Cells were grown at 180 rpm, 37°C to an OD600 of 0.6 then induced with 0.5 mM IPTG and grown overnight at 180 rpm, 18°C. The overnight culture was divided into 5 x 1-mL aliquots for fed and 15 min, 1 h, 2 h, and 3 h glucose deprivation timepoints. Aliquots were pelleted for 5 min at 4000 x g. Fed, 15 min, 1 h, and 2 h cells were resuspended in fed media (phosphate buffered saline, PBS, + 28 mM glucose) while 3 h cells were resuspended in deprivation media (PBS). All cultures were shaken at 180 rpm, 37°C between steps. At 2 h, 1 h, and 15 min before flow analysis cells were pelleted as before and resuspended in deprivation media then returned to shaking at 37°C. On the flow cytometer cells were subjected to λ_ex_ = 405 nm and 485 nm and λ_em_ = 514 nm.

### Flow Cytometry Refeeding Experiments

All assays were performed with the BD FACSAri III Cell Sorter. All sensors and control proteins were expressed in Rosetta(DE3) PanZ was used as a non-fluorescent control, cpGFP was used as a fluorescent control, and PancACe was the experimental condition. Cultures were grown in LB with kanamycin overnight at 37°C then used to induce new cultures. Cells were grown at 180 rpm, 37°C to an OD600 of 0.6 then induced with 0.5 mM IPTG and grown overnight at 180 rpm, 18°C. The overnight culture was divided into 5 x 1-mL aliquots for deprived cells and 15 min, 1h, 2h, and 3h refeeding timepoints. Aliquots were pelleted for 5 min at 4000 x g. Cells were resuspended in deprivation media (PBS) and shaken at 180 rpm, 37°C for 3 h before being moved into fed media (PBS + 28 mM glucose) for the duration of the indicated refeeding time points. Cells waiting to be starved were kept in fed media. Cells were shaken at 37°C between steps. On the flow cytometer cells were subjected to λ_ex_ = 405 nm and 485 nm and λ_em_ = 514 nm.

### Flow Cytometry Data Analysis

Analysis was done in Flowing software. First, the PanZ histogram was used as the negative control to exclude non-fluorescent cells. The region with more fluorescence than PanZ was marked as Region 1. For PancACe and cpGFP the ratio of 485/405 was calculated on a cell by cell basis. This ratio was plotted in a histogram including only cells from region 1. The median of this ratio for each time point was taken. For the deprivation assays, the percent difference was calculated between each deprivation timepoint and the fed control. For the refeeding assays, the percent difference was calculated between each refeeding timepoint and the deprived control.

### Plate Reader Refeeding Experiments

All assays were performed with a BioTek Synergy Neo2 Hybrid Multimode Reader with injectors. All sensors and control proteins were expressed in Rosetta(DE3) PanZ was used as a non-fluorescent control, cpGFP was used as a fluorescent control, and PancACe was the experimental condition. Cultures were grown in LB with kanamycin overnight at 37°C then used to induce new cultures. Cells were grown at 37°C to an OD600 of 0.6 then induced with 0.5 mM IPTG and grown overnight at 18°C. Cultures were split into 2 x 600 µL aliquots and pelleted at 4000 x g for 5 min. The plates were subjected to λ_ex_ = 405 nm and 485 nm and λ_em_ = 514 nm at the indicated read intervals. The 485/405 ratio was calculated, and then the data were normalized to the signal prior to the refeeding ((F_1_-F_0_)/F_0_).

#### Glucose Refeeding

The supernatant was discarded, and the pellets were resuspended in 600 µL either deprivation buffer (100 mM sodium phosphate, 2 mM MgCl_2_, 15 mM (NH_4_)_2_SO_4_ + 0 mM glucose) or refeeding buffer (100 mM sodium phosphate, 2 mM MgCl_2_, 15 mM (NH_4_)_2_SO_4_ + 28 mM glucose). 85 µL of the resuspended cultures were added to 3 wells of a 96-well plate for each sample. Plates were shaken at room temperature for 3 h. Plates were read for 10 min every 30 s with 10 s of shaking between reads. 15 µL of deprivation buffer was added to the fed cells and 15 µL of refeeding buffer was added to deprived cells via the injectors. Plates were read for 60 min every 30 s with 10 s of shaking between reads.

#### Glucose Refeeding +/- 2-Deoxyglucose (2-DG)

The supernatant was discarded, and the pellets were resuspended in 600 µL deprivation buffer (100 mM sodium phosphate, 2 mM MgCl_2_, 15 mM (NH_4_)_2_SO_4_ +/- 28 mM 2-DG). 85 µL of the resuspended cultures were added to 3 wells of a 96-well plate for each sample. Plates were shaken at room temperature for 3 h. Plates were read for 10 min every 30 s with 10 s of shaking between reads. 15 µL of refeeding buffer (100 mM sodium phosphate, 2 mM MgCl_2_, 15 mM (NH_4_)_2_SO_4_ + 28 mM glucose) was added to deprived cells via the injectors. Plates were read for 60 min every 30 s with 10 s of shaking between reads.

#### Acetate Refeeding

The supernatant was discarded, and the pellets were resuspended in 600 µL either deprivation buffer (100 mM sodium phosphate, 2 mM MgCl_2_, 15 mM (NH_4_)_2_SO_4_ + 0 mM glucose) or fed buffer (100 mM sodium phosphate, 2 mM MgCl_2_, 15 mM (NH_4_)_2_SO_4_ + 28 mM glucose). 85 µL of the resuspended cultures were added to 3 wells of a 96-well plate for each sample. Plates were shaken at room temperature for 3 h. Plates were read for 10 min every 30 s with 10 s of shaking between reads. 15 µL of deprivation buffer was added to the fed cells and 15 µL of acetate refeeding buffer (100 mM sodium phosphate, 2 mM MgCl_2_, 15 mM (NH_4_)_2_SO_4_ + 280 µM acetate) was added to deprived cells via the injectors. Plates were read for 60 min every 30 s with 10 s of shaking between reads.

### Human cell handling and preparation of the stable HeLa cell lines

#### General

The 293T and HeLa cells were maintained in standard DMEM with 4.5 g/L glucose (11995-065, Gibco), 10% v/v heat inactivated FBS (10082-147, Gibco), and 1% v/v penicillin/streptomycin (15070063, Gibco) at 37°C and 5% CO_2_. The cells were subcultured by trypsinization (25200-056, Gibco). The cell lines were tested for mycoplasma contamination using the LookOut® Mycoplasma PCR Detection Kit (Sigma Millipore) and were found to be negative for mycoplasma.

#### Lentivirus

293T cells were used to produce lentivirus from the pLJM1 proviral vector containing the PancACe or cpGFP gene with the corresponding localization tags (see sequences below). For each construct, a 10-cm dish of 293T cells in standard DMEM 3% v/v FBS was transfected with 4 µg provirus plasmid (pLJM1 with gene of interest), 4 µg HIV gag-pol plasmid (psPAX2), and 0.57 µg VSV-G plasmid (pMD2.G) using 26 µL of XtremeGene HP (6366244001, Sigma Millipore) in 1 mL of Optimem (11058-021, Gibco). After 24 h, the cells were recovered into 10 mL of standard DMEM with 3% v/v FBS. After 48 h, the virus-containing media was harvested and replaced with 10 mL of standard DMEM with 3% v/v FBS. After 72 h, the virus-containing media was harvested and combined with the media collected at 48 h. The virus was filtered through 0.45 µm to remove cell debris and was stored unconcentrated in 1 mL aliquots at -80 °C. The HeLa cells were transduced in 6-well plates with 1 mL of unconcentrated virus + 10 µg/mL polybrene (TR-1003-G, Sigma Millipore) per well for 24 h. At 48 h post-transduction, the cells were subjected to 1 µg/mL puromycin (J67236.XF, Thermo) to select for infected cells, and the selection was maintained until all the non-transduced HeLa cells were dead (about 72 h). Low passage frozen stocks were prepared of the six resultant polyclonal HeLa cell lines (i.e., PancACe mito, nuc, cyto and cpGFP mito, nuc, cyto) in 10% DMSO-containing standard DMEM (with 10% v/v FBS). For experiments, cells were thawed and passaged for a maximum of 15 passages as sensor/GFP expression and cell viability tended to be compromised after that period.

### Perturbations of the Hela cells

#### General

The PancACe/cpGFP-expressing HeLa cells were counted and plated into glass bottom 96-well plates suitable for confocal microscopy (P96-1.5H-N, CellVis). The control condition (e.g., fed or non-targeting siRNA, “NT”) in each experiment was performed in triplicate wells, and the experimental conditions were performed in singlet wells (up to triplet wells in some experimental replicates). Each experiment was performed at least 3 times on entirely different days and plates (“experimental replicates”). The cells were at a density of approximately 20-40,000 cells per well at the time of imaging. Immediately before beginning a microscopy session, the media was changed to serum and phenol red free DMEM that otherwise matched the contents of the corresponding condition.

#### Deprivation experiments

DMEM containing no glucose, glutamine, pyruvate, phenol red, or FBS (A14430-01 Gibco) was used as the base media. The cells were placed in these media formulations described in **Table S13** for approximately 16 h before imaging.

#### Refeeding experiments

The cells were deprived of glucose as described above (“-glucose” condition). After 16 h, the media was changed to DMEM with 4 mM glutamine and 4.5 g/L glucose but without phenol red or FBS (to prepare the cells for imaging). Images were collected for wells that had been refed for 1 h or 3 h as indicated.

#### Branch chain amino acid (BCAA) deprivation

Powdered DMEM without glucose, glutamine, isoleucine, leucine, valine, sodium pyruvate, sodium bicarbonate, or phenol red was used as the basis for these media formulations (US Biological D9800-36). The powdered DMEM was reconstituted in water using sodium bicarbonate to adjust the pH and sterile filtered prior to use as described by the manufacturer. The media was supplemented with 4.5 g/L glucose, 4 mM glutamine, 0.8 mM leucine, and 10% FBS to generate the BCAA-deprived media (i.e., no isoleucine or valine). The control media was generated in the same way but to it was also added 0.8 mM isoleucine and 0.8 mM valine. Cells were incubated in either the +BCAA or -BCAA media for 24 h prior to imaging.

#### siRNA experiments

siRNAs were purchased as SMARTPools from Horizon (Dharmacon). The siRNAs were reconstituted as instructed by the manufacturer in 1x siRNA buffer (Dharmacon) in RNase-free water (Fisher) to a concentration of 20 µM. These stock aliquots were stored at -20 °C. Each well of a 96-well plate was transfected with 0.3 µL of Lipofectamine 3000 (L3000001 Thermo) and 50 nM siRNA (final concentration) in 300 µL total of Optimem. After incubation for 5-6 hours, the cells were recovered by changing the media to standard DMEM with 10% FBS and 1% pen/strep. The knockdowns were validated by immunoblot (**Figure S14**) as detailed in the **Immunoblotting** section below. The siRNAs and antibodies are listed in **Tables S14 and S15**, respectively. The cells were imaged 48 h after transfection. Immediately before beginning the microscopy session, the media was changed to phenol red and serum free media containing 4.5 g/L glucose and 4 mM glutamine.

### Fluorescence microscopy data collection

The cells were maintained at 37 °C and 5% CO_2_ during imaging using an Oko Lab stage top incubator. A Leica SP8 White Light Confocal microscope was used for imaging. A Leica 20x dry objective was used. Excitation was performed at 405 and 488 nm, and emission was measured at 515 nm. Both excitation lasers were operated at 6% power. The gain of the PMT detector varied depending on the overall brightness of the cells on a given day such that pixel saturation was avoided. The scan parameters were set to 200x scan speed, 2.00 zoom factor, 3.5 µm pinhole. For each well, 10 FOVs were collected.

### Fluorescence microscopy data analysis

The images were processed in a workflow using Fiji. A custom python script was created for high-throughput batch processing of images. The coding of this script was performed with some troubleshooting assistance from ChatGPT versions 3.5 and 4 (OpenAI). The script extracted image metadata and queued two-channel pairs of image files into parallel, non-graphical instances of Fiji running custom ImageJ macros provided by the python script. These macros obtained values for a number of combinations of processing techniques performed on each pair in order to enable later sensitivity analysis and method exploration. One Fiji workflow was selected prior to examination of the full results and was used to report the data in this paper. It performed background subtraction with a sliding rolling ball method of 50 px on each single-channel image. It then created two new images: one with each pixel value as a ratio of the two channels (488 / 405, named as ch01 / ch00) and another with each pixel value as the sum of the two channels. It then created a selection on the sum image using the Otsu auto-threshold algorithm. This sum image was chosen as the basis for the selection as a compromise technique in order to include pixels that had either a large return from one of the channels or a substantial return from both. That selection was then applied to the ratio image. The macro measured and exported desired statistics for the included ratio image pixels, including mean pixel value and total pixel area in the selection. The code is provided in the Supplementary Materials.

Below are equivalent actions that can be done by hand in the Fiji GUI in order to replicate it. For each single-channel image, open the file and select the image window, then perform: “Image” → “Type” → “16-bit”.

“Process → “Subtract background” → “Rolling ball radius: 50 px width” with “Sliding paraboloid” selected).

Process → “Filters” → “Gaussian blur: sigma 2.0”.

“Image Calculator”→ “Divide” and “32-bit float result” selected, [exp wavelength image] / [control wavelength image]. This creates the ratio image in a new window.

“Process” → “Image Calculator” → “Add” and “32-bit float result” selected, again using the two single-channel images. This creates the sum image in a new window.

(With sum image focused) “Process” → “Threshold” → “Otsu” with “Dark background” selected. “Edit → “Selection” → “Create Selection”.

(Focus ratio image window)

“Edit → “Selection” → “Restore Selection”.

“Results” → “Measure”

The parent python script took the one-line exported .csv files compiled them into large .csv files, and then prepared and inserted the .csv data into a local SQLite database file. A view in the SQL database compiled the ∼10 FOVs from each well and created summary statistics. The chosen processing method for reporting data in this paper involved finding means and standard deviations for each well that were both weighted by the number of pixels included in each ratio image’s thresholded selection. The weighted summary statistics were then recalculated after excluding FOVs whose mean pixel values were three or more area-weighted standard deviations from the area-weighted mean of the well. The code is provided in the Supplementary Materials.

A further series of views and tables was then created in the SQLite database to enable exploration and comparison of processing methodologies as well as sensitivity analysis. For the purposes of compiling and reporting data from the prior-chosen analytic workflow, well-level data was exported from the SQLite database in .csv format and then processed manually in MS Excel and Graphpad Prism as follows.

The PancACe measurement was divided by the corresponding cpGFP measurement for a given condition (i.e., PancACe/cpGFP). At this stage, the data were subjected to statistical testing. For visualizing data, they were further normalized by dividing the experimental by the corresponding control condition (e.g., Fed/deprived) to yield a fold change. For plots, the percentage change is displayed from the following calculation: -(1-(foldchange))*100. The statistical analyses are described in the respective captions and in the **Data Handling and Statistics** section below.

### PicoProbe^TM^ fluorometric acetyl-CoA assay

A 10-cm plate (∼2x10^6^ cells) of HeLa cells was given complete media (fed) or media lacking glucose or glutamine for 16 h. The media was removed, and the plates were washed with 2 mL of 1x PBS pH 7.4 (10010023, Gibco). The cells were treated with 1 mL of trypsin, incubated for five minutes at 37°C, and then pelleted for 5 min at 200 x g at 4°C. For normalization, the cell density of each plate was determined using a Countess 3 (Thermofisher). The supernatant was discarded, and the cells were resuspended in 50 µL of “MS buffer” (210 mM mannitol, 70 mM sucrose, 5 mM Tris-HCl, and 1 mM EDTA, pH 7.5). 10 µL of 1 M perchloric acid (Sigma Millipore) was added to resuspended cells and sonicated at 10% power for three one-second cycles with three-second rests between cycles. The lysates were pelleted at 17,000 x g at 4°C for 10 minutes. 3 µL of 3 M potassium bicarbonate was added to each clarified lysate and mixed before incubating on ice for 5 min. The lysate was pelleted at 17,000 x g at 4°C for 1 min. The PicoProbe^TM^ assay (Sigma Millipore) and analysis were performed according to the manufacturer’s protocols using the picomole standard curve (n = 1). The only adaptation was for the blank; a 1:5 dilution was prepared for the fluorescent probe in the acetyl-CoA assay buffer.

### Immunoblotting

HeLa cell samples were treated in the same manner as the corresponding microscopy experiment. The only difference was that conditions for immunoblotting were scaled up to a 6-well plate format. The cells were washed with cold 1x PBS (10010023, Gibco) and then harvested in cold 1x PBS. The Epiquik Histone Extraction Kit (Epigentek) was used for lysis and histone extraction as directed by the manufacturer. Briefly, the cell pellets were resuspended in 100 µL of 1x Pre-Lysis buffer and lysed on ice for 10 min. The cells were pelleted at high speed for 1 min, and the supernatant was reserved (“lysate” or “total soluble fraction”). The pellet was resuspended in 30 µL of the Lysis buffer and incubated on ice for 30 min. The material was pelleted, and the supernatant was reserved (“histones”) and neutralized with 9 µL of Balance buffer as directed. SDS loading buffer was added to both lysate and histone samples for immunoblotting.

The protein samples were separated by SDS-PAGE gel (either 12% Bis Tris or 4-12% Bis Tris) in 1x MES and transferred to a PVDF membrane by semi-dry transfer in Towbin buffer (20 V for 30 min). For the p300 and ACC blots, the proteins were separated on a 4-12% Bis Tris gel in 1x MOPS for 1.5 h and were transferred for 45 min at 20 V. The membranes were blocked with 3% or 5% milk according to the antibody manufacturer’s instructions for 1 h at room temperature. The membranes were incubated with primary antibodies in TBST with 3% BSA at 4°C overnight and then washed with TBST 3x. The membranes were incubated with secondary antibodies (Licor) in TBST for 1 h at room temperature and then washed with TBST 3x and 1x PBS 1x before imaging using an Odyssey Imager. Densitometry was performed within the Licor Image Studio software.

### Data Handling and Statistics

Microsoft Excel was used for initial data processing. SQLite was used for initial processing of the image data from Fiji as described above. GraphPad Prism was used to make all of the plots and for fitting data. The number of replicates is given in each figure and/or figure caption. The data are displayed with the standard error of the mean (SEM) or the standard deviations (SD) as indicated in the respective figure captions. In general, the SEM was used for the biochemical assays, and SD was used for the cell-based assays. Data fitting (linear and non-linear models) was performed in GraphPad Prism. Outlier testing was performed in GraphPad Prism using the Grubbs’ method. The statistical testing for the cell-based experiments was performed in GraphPad Prism. Paired two-tailed t tests, two-way ANOVA, and/or Tukey’s multiple comparison tests were used, and each figure caption indicates the specific test that was used. We note that for the HeLa cell data, we performed paired two-tailed t tests on data that was not normalized to the control condition when we were making a comparison between an experimental and control condition. All of the p values are shown in **Tables S1-S9** and/or the respective figure caption.

## Acknowledgments

We acknowledge the Cell Imaging Core at the University of Utah for use of equipment (Leica SP8 Confocal Microscope) and thank Dr. Michael Bridge for his assistance. We also acknowledge the DNA/Peptide Synthesis, Flow Cytometry, and Drug Discovery Cores at the University of Utah. We acknowledge ChapGPT versions 3.5 and 4 (OpenAI) for assistance in writing the Python script for the image processing. We acknowledge Drs. Kathryn Wellen and Gregory Ducker for helpful discussions.

## Funding

National Institutes of Health grant R35GM143080 (KLD)

University of Utah Undergraduate Research Opportunity Program (GAD)

## Author contributions

Conceptualization: JJS, TRV, KLD

Methodology: JJS, TRV, AHA, RB, DME, KLD

Software: ARC

Formal analysis: JJS, AHA, RB, DME, KLD

Investigation, JJS, TRV, AHA, RB, DME, GAD, KLD

Visualization: JJS, RB, DME, KLD

Writing- original draft: JJS, KLD

Writing- reviewing and editing: JJS, TRV, AHA, RB, ARC, DME, KLD

Supervision: KLD

JJS, TRV, and KLD designed the experiments. JJS performed the cloning, protein purification, biochemical assays with the sensors, and the E. coli assays. TRV generated the stable cell lines and established the microscopy assay in the HeLa cells. AHA performed the PicoProbe assay, some cloning and protein purification, and the linker composition experiments. RB performed the molecular modeling of PanZ and PancACe. ARC wrote and employed the Python, Fiji macro, and SQLite scripting codes for image data acquisition and initial processing of data to the well level. DME performed the surface plasmon resonance (SPR) assays. GAD performed some cloning and protein purification and the YFP and BFP experiments. KLD performed the immunoblot assays, the microscopy assays in the HeLa cells, and the analysis of the image data. JJS and KLD wrote the manuscript with input from all authors.

## Data and materials availability

All data are available in the main text or the supplementary materials. A description of the code used for batch image processing is provided in the supplementary materials. The plasmids described herein are available upon request via a materials transfer agreement (MTA). Certain plasmids (Ecoli PancACe, Ecoli cpGFP, Nuclear PancACe, Nuclear cpGFP, Cytoplasmic PancACe, Cytoplasmic cpGFP, Mitochondrial PancACe, and Mitochondrial cpGFP) are available on Addgene (identifiers pending).

