## Supplemental materials for "A genetically-encoded fluorescent biosensor for visualization of acetyl-CoA in live cells"

Joseph J. Smith *et al.*

**This PDF file includes:**

Supplementary Text
Figs. S1 to S17
Tables S1 to S15
Data Files S1 to S7

**Supplementary Text**

Amino acid sequences for the proteins

CFP-PanZ
MHHHHHHENLYFQSDLYDDDDKDPMVSKGEELFTGVVPILVELDGDVNGHRFSVSGEGEGDA
TYGKLTLKFICTTGKLPVPWPTLVTTLTWGVQCFSRYPDHMKQHDFFKSAMPEGYVQERTIFF
KDDGNYKTRAEVKFEGDTLVNRIELKGIDFKEDGNILGHKLEYNYISHNVYITADKQKNGIKAHF
KIRHNIEDGSVQLADHYQQNTPIGDGPVLLPDNHYLSTQSALSKDPNEKRDHMVLLEFVTAAR
MLPVDLMKLTIIRLEKFSDQDRIDLQKIWPEYSPSSLQVDDNHRIYAARFNERLLAAVRVTLSGT
EGALDSLRVREVTRRRGVGQYLLEEVLRNNPGVSCWWMADAGVEDRGVMTAFMQALGFTA
QQGGWEKC*

HTSD8.05 6xHis-TEV-PancACe
Key: PanZ, cpGFP
MHHHHHHENLYFQSMKLTIIRLEKFSDQDRIDLQKIWPEYSPSSLQVDDNHRIYAARFNERLLA
AVRVTLSGTEGALDSLRVRGANVYIKADKQKNGIKANFKIRHNIEDGGVQLAYHYQQNTPIGDG
PVLLPDNHYLSVQSKLSKDPNEKRDHMVLLEFVTAAGITLGMDELYKGGTGGSMVSKGEELFT
GVVPILVELDGDVNGHKFSVSGEGEGDATYGKLTLKFICTTGKLPVPWPTLVTTLTYGVQCFSR
YPDHMKQHDFFKSAMPEGYIQERTIFFKDDGNYKTRAEVKFEGDTLVNRIELKGIDFKEDGNIL
GHKLEYNGAEVTRRRGVGQYLLEEVLRNNPGVSCWWMADAGVEDRGVMTAFMQALGFTAQ
QGGWEKC*

HTSD1.00 6xHis-TEV-cpGFP
(Plasmid name: Ecoli cpGFP)
MHHHHHHENLYFQSNVYIKADKQKNGIKANFKIRHNIEDGGVQLAYHYQQNTPIGDGPVLLPDN
HYLSVQSKLSKDPNEKRDHMVLLEFVTAAGITLGMDELYKGGTGGSMVSKGEELFTGVVPILV
ELDGDVNGHKFSVSGEGEGDATYGKLTLKFICTTGKLPVPWPTLVTTLTYGVQCFSRYPDHMK
QHDFFKSAMPEGYIQERTIFFKDDGNYKTRAEVKFEGDTLVNRIELKGIDFKEDGNILGHKLEYN
*

*Position Screen*
cpGFP plus linkers insert
ASNVYIKADKQKNGIKANFKIRHNIEDGGVQLAYHYQQNTPIGDGPVLLPDNHYLSVQSKLSKD
PNEKRDHMVLLEFVTAAGITLGMDELYKGGTGGSMVSKGEELFTGVVPILVELDGDVNGHKFS
VSGEGEGDATYGKLTLKFICTTGKLPVPWPTLVTTLTYGVQCFSRYPDHMKQHDFFKSAMPEG
YIQERTIFFKDDGNYKTRAEVKFEGDTLVNRIELKGIDFKEDGNILGHKLEYNAS

Positions of different constructs where cpGFP plus linkers insert was used
MHHHHHHENLYFQSMKLTIIRLEKFSDQDRIDLQKIW(HTSD6.00)PEY(HTSD6.01)SPSSL(HTS
D6.02)QVDDNHRIYAARFN(HTSD6.03)ERLLAAVRVTLSGTEGALDSLRVR(HTSD6.04)EV(HT
SD6.05)TRRRG(HTSD6.06)VGQYLLEEVLRNNP(HTSD6.07)GVSCWWMAD(HTSD6.08)AG(H
TSD6.09)VED(HTSD6.10)RGVMTAFMQALG(HTSD6.11)FTAQQGGWEKC*

Example HTSD6.04
MHHHHHHENLYFQSMKLTIIRLEKFSDQDRIDLQKIWPEYSPSSLQVDDNHRIYAARFNERLLA
AVRVTLSGTEGALDSLRVRASNVYIKADKQKNGIKANFKIRHNIEDGGVQLAYHYQQNTPIGDG
PVLLPDNHYLSVQSKLSKDPNEKRDHMVLLEFVTAAGITLGMDELYKGGTGGSMVSKGEELFT
GVVPILVELDGDVNGHKFSVSGEGEGDATYGKLTLKFICTTGKLPVPWPTLVTTLTYGVQCFSR
YPDHMKQHDFFKSAMPEGYIQERTIFFKDDGNYKTRAEVKFEGDTLVNRIELKGIDFKEDGNIL

GHKLEYNASEVTRRRGVGQYLLEEVLRNNPGVSCWWMADAGVEDRGVMTAFMQALGFTAQ
QGGWEKC*

*Linker Length Screen*
HTSD8.00_MHHHHHHENLYFQSMKLT…PanZ…LVRGASNVY…cpGFP…EYNGASEVT…PanZ…EKC*
HTSD8.01_MHHHHHHENLYFQSMKLT…PanZ…LVRGASNVY…cpGFP…EYNGAEVT…PanZ…EKC*
HTSD8.02_MHHHHHHENLYFQSMKLT…PanZ…LVRGASNVY…cpGFP…EYNGEVT…PanZ…EKC*
HTSD8.03_MHHHHHHENLYFQSMKLT…PanZ…LVRGASNVY…cpGFP…EYNEVT…PanZ…EKC*
HTSD8.04_MHHHHHHENLYFQSMKLT…PanZ…LVRGANVY…cpGFP…EYNGASEVT…PanZ…EKC*
HTSD8.05_MHHHHHHENLYFQSMKLT…PanZ…LVRGANVY…cpGFP…EYNGAEVT…PanZ…EKC*
HTSD8.06_MHHHHHHENLYFQSMKLT…PanZ…LVRGANVY…cpGFP…EYNGEVT…PanZ…EKC*
HTSD8.07_MHHHHHHENLYFQSMKLT…PanZ…LVRGANVY…cpGFP…EYNEVT…PanZ…EKC*
HTSD8.08_MHHHHHHENLYFQSMKLT…PanZ…LVRGNVY…cpGFP…EYNGASEVT…PanZ…EKC*
HTSD8.09_MHHHHHHENLYFQSMKLT…PanZ…LVRGNVY…cpGFP…EYNGAEVT…PanZ…EKC*
HTSD8.10_MHHHHHHENLYFQSMKLT…PanZ…LVRGNVY…cpGFP…EYNGEVT…PanZ…EKC*
HTSD8.11_MHHHHHHENLYFQSMKLT…PanZ…LVRGNVY…cpGFP…EYNEVT…PanZ…EKC*
HTSD8.12_MHHHHHHENLYFQSMKLT…PanZ…LVRNVY…cpGFP…EYNGASEVT…PanZ…EKC*
HTSD8.13_MHHHHHHENLYFQSMKLT…PanZ…LVRNVY…cpGFP…EYNGAEVT…PanZ…EKC*
HTSD8.14_MHHHHHHENLYFQSMKLT…PanZ…LVRNVY…cpGFP…EYNGEVT…PanZ…EKC*
HTSD8.15_MHHHHHHENLYFQSMKLT…PanZ…LVRNVY…cpGFP…EYNEVT…PanZ…EKC*

Example HTSD8.05(PancACe)
(Plasmid name: Ecoli PancACe)
MHHHHHHENLYFQSMKLTIIRLEKFSDQDRIDLQKIWPEYSPSSLQVDDNHRIYAARFNERLLA
AVRVTLSGTEGALDSLRVRGANVYIKADKQKNGIKANFKIRHNIEDGGVQLAYHYQQNTPIGDG
PVLLPDNHYLSVQSKLSKDPNEKRDHMVLLEFVTAAGITLGMDELYKGGTGGSMVSKGEELFT
GVVPILVELDGDVNGHKFSVSGEGEGDATYGKLTLKFICTTGKLPVPWPTLVTTLTYGVQCFSR
YPDHMKQHDFFKSAMPEGYIQERTIFFKDDGNYKTRAEVKFEGDTLVNRIELKGIDFKEDGNIL
GHKLEYNGAEVTRRRGVGQYLLEEVLRNNPGVSCWWMADAGVEDRGVMTAFMQALGFTAQ
QGGWEKC*

*Linker Composition Screen*
MHHHHHHENLYFQSMKLT…PanZ…LVRXYNVY…cpGFP…EYNXYEVT…PanZ…EKC*
Where XY = GG, GE, EG, EE QQ, RR, or KK

*cpYFP and cpBFP variants*

HTSD1.01 6xHistidine TEV cpYFP
MHHHHHHENLYFQSNVYIKADKQKNGIKANFKIRHNIEDGGVQLAYHYQQNTPIGDGPVLLPDN
HYLSYQSVLSKDPNEKRDHMVLLEFVTAAGITLGMDELYKGGTGGSMVSKGEELFTGVVPILV
ELDGDVNGHKFSVSGEGEGDATYGKLTLKFICTTGKLPVPWPTLVTTLTYGVQCFSRYPDHMK
QHDFFKSAMPEGYIQERTIFFKDDGNYKTRAEVKFEGDTLVNRIELKGIDFKEDGNILGHKLEYN
*

HTSD1.02 6xHistidine TEV cpBFP
MHHHHHHENLYFQSNVYIKADKQKNGIKANFKIRHNIEDGGVQLAYHYQQNTPIGDGPVLLPDN
HYLSVQSKLSKDPNEKRDHMVLLEFVTAAGITLGMDELYKGGTGGSMVSKGEELFTGVVPILV
ELDGDVNGHKFSVSGEGEGDATYGKLTLKFICTTGKLPVPWPTLVTTLSHGVQCFSRYPDHMK
QHDFFKSAMPEGYIQERTIFFKDDGNYKTRAEVKFEGDTLVNRIELKGIDFKEDGNILGHKLEYN
*

*Sequences in the pLJM1 plasmid (lentiviral expression)*
Key: FLAG and HA tags are underlined; **localization tags**(*50*) **are bolded**
Nuclear PancACe
MVTG**PKKKRKV**DYKDDDDKLDGGYPYDVPDYAARGYQTSLYKKAGSTMGHMKLTIIRLEKFS
DQDRIDLQKIWPEYSPSSLQVDDNHRIYAARFNERLLAAVRVTLSGTEGALDSLRVRGANVYIK
ADKQKNGIKANFKIRHNIEDGGVQLAYHYQQNTPIGDGPVLLPDNHYLSVQSKLSKDPNEKRD
HMVLLEFVTAAGITLGMDELYKGGTGGSMVSKGEELFTGVVPILVELDGDVNGHKFSVSGEGE
GDATYGKLTLKFICTTGKLPVPWPTLVTTLTYGVQCFSRYPDHMKQHDFFKSAMPEGYIQERTI
FFKDDGNYKTRAEVKFEGDTLVNRIELKGIDFKEDGNILGHKLEYNGAEVTRRRGVGQYLLEEV
LRNNPGVSCWWMADAGVEDRGVMTAFMQALGFTAQQGGWEKC*

Nuclear cpGFP
MVTG**PKKKRKV**DYKDDDDKLDGGYPYDVPDYAARGYQTSLYKKAGSTMGHNVYIKADKQKN
GIKANFKIRHNIEDGGVQLAYHYQQNTPIGDGPVLLPDNHYLSVQSKLSKDPNEKRDHMVLLEF
VTAAGITLGMDELYKGGTGGSMVSKGEELFTGVVPILVELDGDVNGHKFSVSGEGEGDATYGK
LTLKFICTTGKLPVPWPTLVTTLTYGVQCFSRYPDHMKQHDFFKSAMPEGYIQERTIFFKDDGN
YKTRAEVKFEGDTLVNRIELKGIDFKEDGNILGHKLEYN*

Cytoplasmic PancACe
MVTG**LQKKLEELELD**DYKDDDDKLDGGYPYDVPDYAARGYQTSLYKKAGSTMGHMKLTIIRLE
KFSDQDRIDLQKIWPEYSPSSLQVDDNHRIYAARFNERLLAAVRVTLSGTEGALDSLRVRGANV
YIKADKQKNGIKANFKIRHNIEDGGVQLAYHYQQNTPIGDGPVLLPDNHYLSVQSKLSKDPNEK
RDHMVLLEFVTAAGITLGMDELYKGGTGGSMVSKGEELFTGVVPILVELDGDVNGHKFSVSGE
GEGDATYGKLTLKFICTTGKLPVPWPTLVTTLTYGVQCFSRYPDHMKQHDFFKSAMPEGYIQE
RTIFFKDDGNYKTRAEVKFEGDTLVNRIELKGIDFKEDGNILGHKLEYNGAEVTRRRGVGQYLL
EEVLRNNPGVSCWWMADAGVEDRGVMTAFMQALGFTAQQGGWEKC*

Cytoplasmic cpGFP
MVTG**LQKKLEELELD**DYKDDDDKLDGGYPYDVPDYAARGYQTSLYKKAGSTMGHNVYIKADK
QKNGIKANFKIRHNIEDGGVQLAYHYQQNTPIGDGPVLLPDNHYLSVQSKLSKDPNEKRDHMV
LLEFVTAAGITLGMDELYKGGTGGSMVSKGEELFTGVVPILVELDGDVNGHKFSVSGEGEGDA
TYGKLTLKFICTTGKLPVPWPTLVTTLTYGVQCFSRYPDHMKQHDFFKSAMPEGYIQERTIFFK
DDGNYKTRAEVKFEGDTLVNRIELKGIDFKEDGNILGHKLEYN*

Mitochondrial PancACe
**MVLATRVFSLVGKRAISTSVCVRAHT**GDYKDDDDKLDGGYPYDVPDYAARGYQTSLYKKAGS
TMGHMKLTIIRLEKFSDQDRIDLQKIWPEYSPSSLQVDDNHRIYAARFNERLLAAVRVTLSGTEG

ALDSLRVRGANVYIKADKQKNGIKANFKIRHNIEDGGVQLAYHYQQNTPIGDGPVLLPDNHYLS
VQSKLSKDPNEKRDHMVLLEFVTAAGITLGMDELYKGGTGGSMVSKGEELFTGVVPILVELDG
DVNGHKFSVSGEGEGDATYGKLTLKFICTTGKLPVPWPTLVTTLTYGVQCFSRYPDHMKQHDF
FKSAMPEGYIQERTIFFKDDGNYKTRAEVKFEGDTLVNRIELKGIDFKEDGNILGHKLEYNGAEV
TRRRGVGQYLLEEVLRNNPGVSCWWMADAGVEDRGVMTAFMQALGFTAQQGGWEKC*

Mitochondrial cpGFP
**MVLATRVFSLVGKRAISTSVCVRAHT**GDYKDDDDKLDGGYPYDVPDYAARGYQTSLYKKAGS
TMGHNVYIKADKQKNGIKANFKIRHNIEDGGVQLAYHYQQNTPIGDGPVLLPDNHYLSVQSKLS
KDPNEKRDHMVLLEFVTAAGITLGMDELYKGGTGGSMVSKGEELFTGVVPILVELDGDVNGHK
FSVSGEGEGDATYGKLTLKFICTTGKLPVPWPTLVTTLTYGVQCFSRYPDHMKQHDFFKSAMP
EGYIQERTIFFKDDGNYKTRAEVKFEGDTLVNRIELKGIDFKEDGNILGHKLEYN*

Python script for batch image processing in Fiji

A reconstruction of some of the key commands used by the macro for the chosen method is provided below, with relevant values substituted for some variables or lists of variable values that appear in the original macro script. Image metadata supplied to the macro from the parent python script and processing technique metadata generated by the macro for each measurement were also exported along with each measurement result using the setResult() command.
The specifics of the metadata exporting are omitted here for brevity.

```
run("Set Measurements...", "area area_fraction mean standard min redirect=None decimal=3");
run("Input/Output...", "jpeg=85 gif=-1 file=.csv use_file save_column");
rolling_ball_size = 50
gb_sigma = 2
thr_method = "Otsu"
function setMeasure() {
	meas_count = meas_count + 1;
	run("Measure");
	setResult("[first_metadata_var_name]", 0, [value]);
	setResult("[fsecond_metadata_var_name]", 0, [value]);
	….
	print("Saving results for string " + meas_str);
	saveAs("Results", pair_csv_base_fullpath + toString(meas_count) + ".csv");
}
open([ch00_raw_filepath]);
ch00_title = getInfo("image.title");
open([ch01_raw_filepath]);
ch01_title = getInfo("image.title");
selectWindow(ch00_title);
run("16-bit");
run("Subtract Background...", "rolling=" + toString(rolling_ball_size) + " sliding");
selectWindow(ch01_title);
run("16-bit");
run("Subtract Background...", "rolling=" + toString(rolling_ball_size) + " sliding");
 selectWindow(ch00_title);
run("Gaussian Blur...", "sigma="+gb_sigma);
selectWindow(ch01_title);
```

```
run("Gaussian Blur...", "sigma="+gb_sigma);
 imageCalculator("Divide create 32-bit", ch01_title, ch00_title);
 rename("ratio_gb");
 ratio_title = getInfo("image.title");
 imageCalculator("Add create 32-bit", ch01_title, ch00_title);
rename("sum_gb");
sum_title = getInfo("image.title");
selectWindow(sum_title);
setAutoThreshold(thr_method + " dark");
run("Create Selection");
selectWindow(ratio_title);
run("Restore Selection");
setMeasure();
```

View SQL code

Below is the view SQL code, which includes use of the “stats” extension for SQLite (maintained by Anton Zhiyanov, https://github.com/nalgeon/sqlean/blob/main/docs/stats.md).

```
select *,
sum(pixval_mean * Area) FILTER (where abs(pos_aw_zscore) < 3) OVER (partition by
m_well_id)
/
sum(Area) FILTER (where abs(pos_aw_zscore) < 3) OVER (partition by m_well_id)
as well_aw_mean_no
from
(
select m_well_id, macsr_id, pkid, meas_str_id, pixval_mean, well_aw_mean, well_uw_mean,
Area, well_area_all
--, stddev_samp(pixval_mean)
,SQRT(
SUM(Area * power((pixval_mean - well_aw_mean),2)
) over w2
/(
(
CAST(COUNT(*) over w2 - 1 AS REAL) /
CAST(COUNT(*) over w2 AS REAL)
) *
SUM(Area) over w2
)
) AS well_aw_stdev
,(pixval_mean - well_aw_mean) /
SQRT(
SUM(Area * power((pixval_mean - well_aw_mean),2)
) over w2
/
(
(
CAST(COUNT(*) over w2 - 1 AS REAL) /
CAST(COUNT(*) over w2 AS REAL)
) *
```

```
SUM(Area) over w2
)
)
as pos_aw_zscore
, sqrt(sum(sq_diff_uw) over w2 / (count(*) over w2 - 1 ) ) as well_uw_stdev
,(pixval_mean - well_uw_mean) / sqrt(sum(sq_diff_uw) over w2 / (count(*) over w2 - 1 ) )
as pos_uw_zscore
from
(
select m_well_id, macsr_id, pkid, meas_str_id, pixval_mean, Area
,avg(pixval_mean) over ww as well_uw_mean
--,stddev_samp(pixval_mean) over ww as well_uw_stdev
,sum(Area) over ww as well_area_all
,sum(pixval_mean * Area) over ww / sum(Area) over ww as well_aw_mean
,avg(pixval_mean) over ww as well_uw_mean
,
power((pixval_mean - avg(pixval_mean) over ww),2)
as sq_diff_uw
,
power((pixval_mean - SUM(pixval_mean * Area) OVER ww / SUM(Area) OVER ww),2)
as sq_diff_aw
from fast_supertable
where targ_channel in ('ratio','quot')
window ww as (partition by m_well_id)
)
window w2 as (partition by m_well_id)
--group by full_well_id
)
```

a

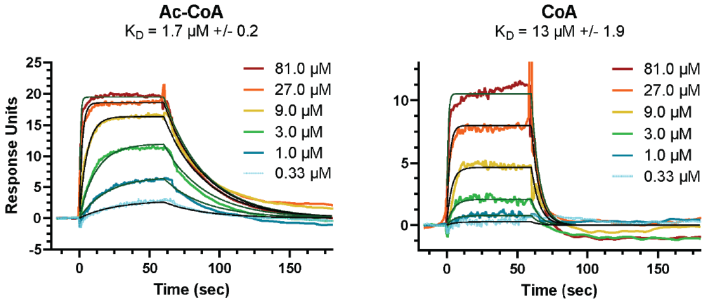


c

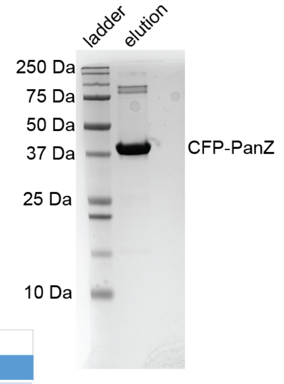


b

| Ac-CoA binding to PanZ-CFP | | | | | |
|---|---|---|---|---|---|
| | Kinetic Fit | | | | |
| | Rmax (RU) | $K_D$ (µM) | $k_{on}$ ($M^{-1}$•$S^{-1}$) | $k_{off}$ ($s^{-1}$) | $t_{1/2;}$ ln2/k (s) |
| Replicate 1 | 20 | 1.98 | 1.69E+04 | 0.034 | 20.7 |
| Replicate 2 | 19 | 1.86 | 1.90E+04 | 0.036 | 19.5 |
| Replicate 3 | 22 | 1.95 | 2.37E+04 | 0.046 | 15.0 |
| Replicate 1 | 32 | 2.03 | 3.05E+04 | 0.062 | 11.2 |
| Replicate 2 | 32 | 0.86 | 8.62E+04 | 0.074 | 9.4 |
| Average | | 1.74 | 3.53E+04 | 0.050 | 15.2 |
| Standard Error | | 0.20 | 1.16E+04 | 0.007 | 2.0 |

| CoA binding to PanZ-CFP | | | | | |
|---|---|---|---|---|---|
| | Kinetic Fit | | | | |
| | Rmax (RU) | $K_D$ (µM) | $k_{on}$ ($M^{-1}$•$S^{-1}$) | $k_{off}$ ($s^{-1}$) | $t_{1/2;}$ ln2/k (s) |
| Replicate 1 | 13 | 15.2 | 1.16E+04 | 0.176 | 3.9 |
| Replicate 2 | 9.1 | 8.6 | 1.48E+04 | 0.127 | 5.5 |
| Replicate 1 | 30 | 18.2 | 1.19E+04 | 0.217 | 3.2 |
| Replicate 2 | 25 | 10.0 | 2.14E+04 | 0.213 | 3.3 |
| Average | | 13.0 | 1.49E+04 | 0.183 | 4.0 |
| Standard Error | | 1.9 | 1.97E+03 | 0.018 | 0.5 |

**Fig. S1. Surface plasmon resonance analysis of CFP-PanZ.** a) Representative plot for CFP-PanZ binding to acetyl-CoA (left) or CoA (right). The dissociation constants (“$K_D$”) for the average of all of the replicates performed are shown in the figure. b) Table of the kinetic data for acetyl-CoA (top) or CoA (bottom) binding to CFP-PanZ. The experimental replicates across two days of data collection are shown. c) Gel image of CFP-PanZ (Coomassie blue staining) analyzed on a 12% Bis Tris gel.

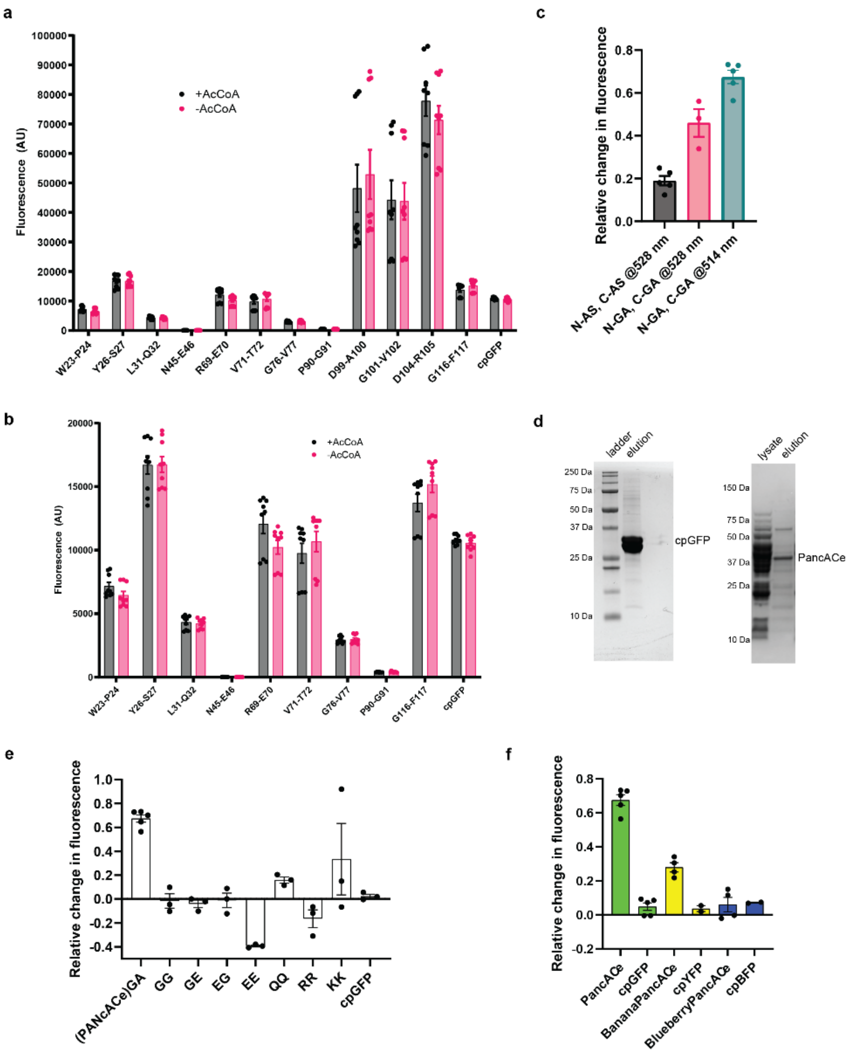


**Fig. S2**. **Engineering of an acetyl-CoA biosensor from PanZ.** a) Fluorescence intensity for the insertion library constructs with and without 1 mM acetyl-CoA, n = 9 (3 sets of technical replicates from 3 experimental replicates), $\lambda_{ex}$ = 485, $\lambda_{em}$ = 528 nm b) Zoom in view of the data in part a. c) Relative fluorescence intensity of the AS and GA linker variants of the R69/E70 insertion site in the presence of 1 mM acetyl-CoA compared to in the absence of acetyl-CoA. The data are replotted from the previous figures (**Figures 1b** and **1c**). Emission wavelength is indicated in panel. d) Representative gel images of PancACe and cpGFP (Coomassie blue staining). PancACe was analyzed on a 4-12% Bis Tris gel, while cpGFP was analyzed on a 12% Bis Tris gel. e) Analysis of the linker composition variants. The data are displayed as relative fluorescence in the presence of 1 mM acetyl-CoA compared to in the absence of acetyl-CoA, $\lambda_{ex}$ = 485, $\lambda_{em}$ = 514 nm. f) Analysis of banana-PancACe ($\lambda_{ex}$ = 505, $\lambda_{em}$ = 535 nm) and blueberry-PancACe ($\lambda_{ex}$ = 380, $\lambda_{em}$ = 440 nm). The data are displayed as relative fluorescence in the presence of 1 mM acetyl-CoA compared to in the absence of acetyl-CoA. For panels c, e, and f, the data are displayed as the relative fluorescence: $(F_1-F_0)/F_0$). n = 3-5, SEM shown.

a

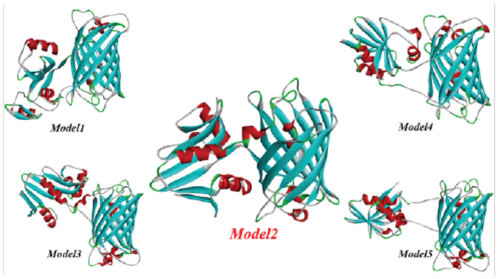


| Model No. | YASARA Energy (kJ/mol) |
|---|---|
| Model 1 | -185882.4 |
| *Model 2* | *-196993.7* |
| Model 3 | -189670.5 |
| Model 4 | -186935.4 |
| Model 5 | -188354.4 |

b

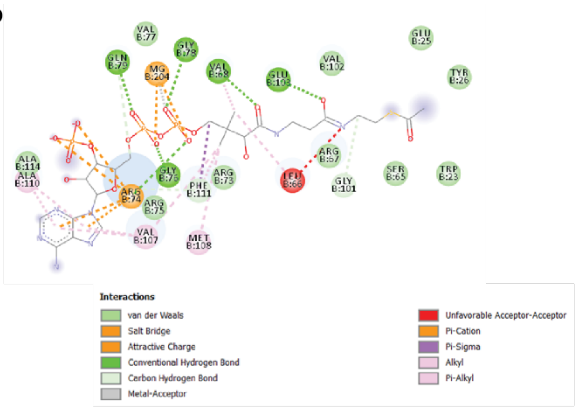


c

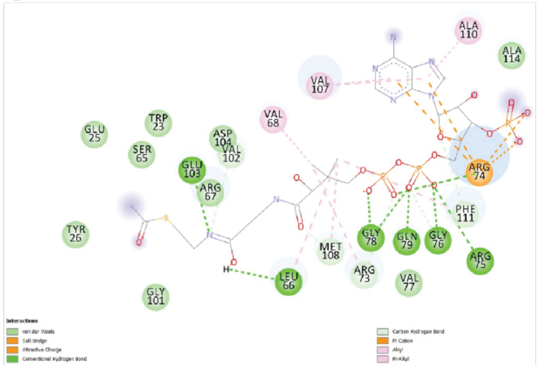

d

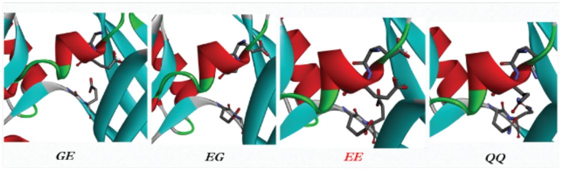


**Fig. S3. Homology modeling and energy minimization of PanZ and PancACe.** a) Models generated using the Robetta server for the PanZ(1-69)–GA-cpGFP-GA-PanZ(70-137) (PancACe) construct. Results of the energy minimization of five models are shown in the table inset. Model 2 is highlighted in red. b) Crystal structure of PanZ (PDB 5lS7) showing binding site interactions with acetyl-CoA. c) PanZ(1-69)–GA-cpGFP-GA-PanZ(70-137) (PancACe) construct docked with Acetyl-CoA displays displacement from previous active site coordinates in space, resulting in reduced arginine anchor interactions and phosphate salt bridge interactions. d) Examination of the linkers of the GE, EG, EE and QQ constructs showed charged glutamate residues entering the cpGFP β-barrel.

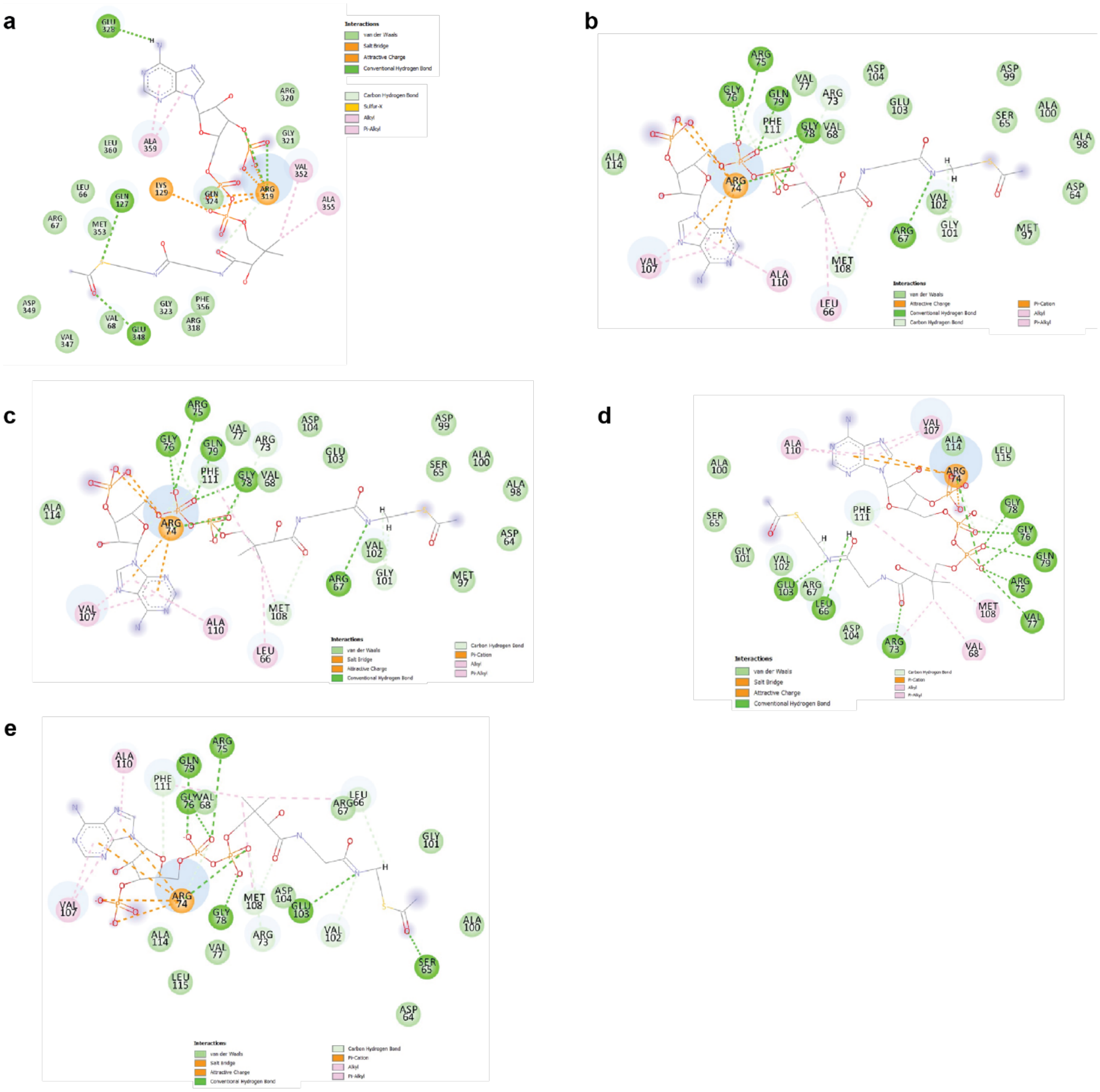


**Fig. S4. Clusters 1-5 of the backdock of acetyl-CoA into PanZ (PDB 5IS7).**

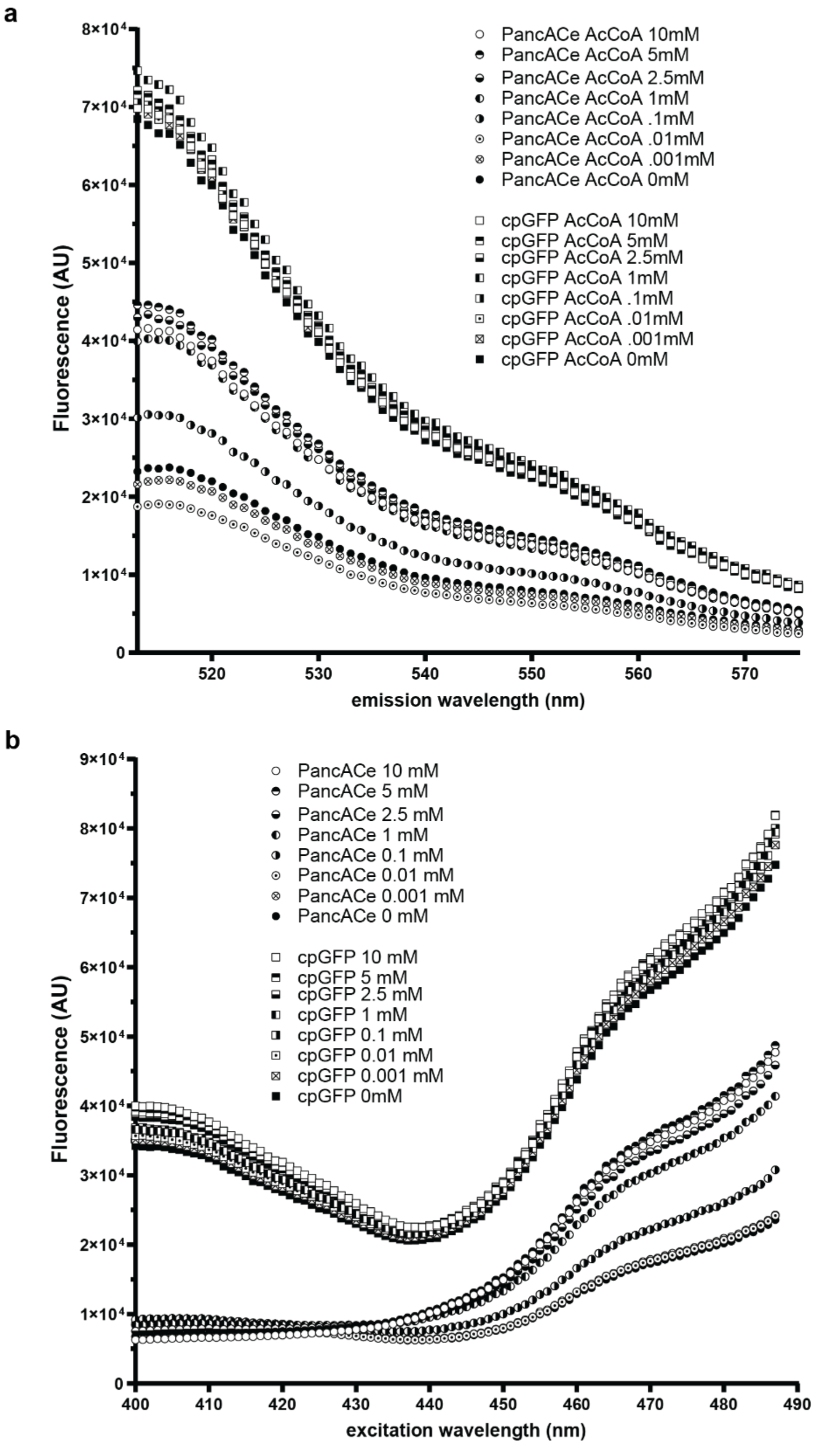


**Fig. S5**. **Fluorescence spectra of PancACe and cpGFP.** a) Average emission spectra of PancACe and cpGFP in the presence of the indicated concentrations of acetyl-CoA, $\lambda_{ex}$ = 485 nm, n = 3. b) Average excitation spectra of PancACe and cpGFP in the presence of the indicated concentrations of acetyl-CoA, $\lambda_{em}$ = 514 nm, n = 3.

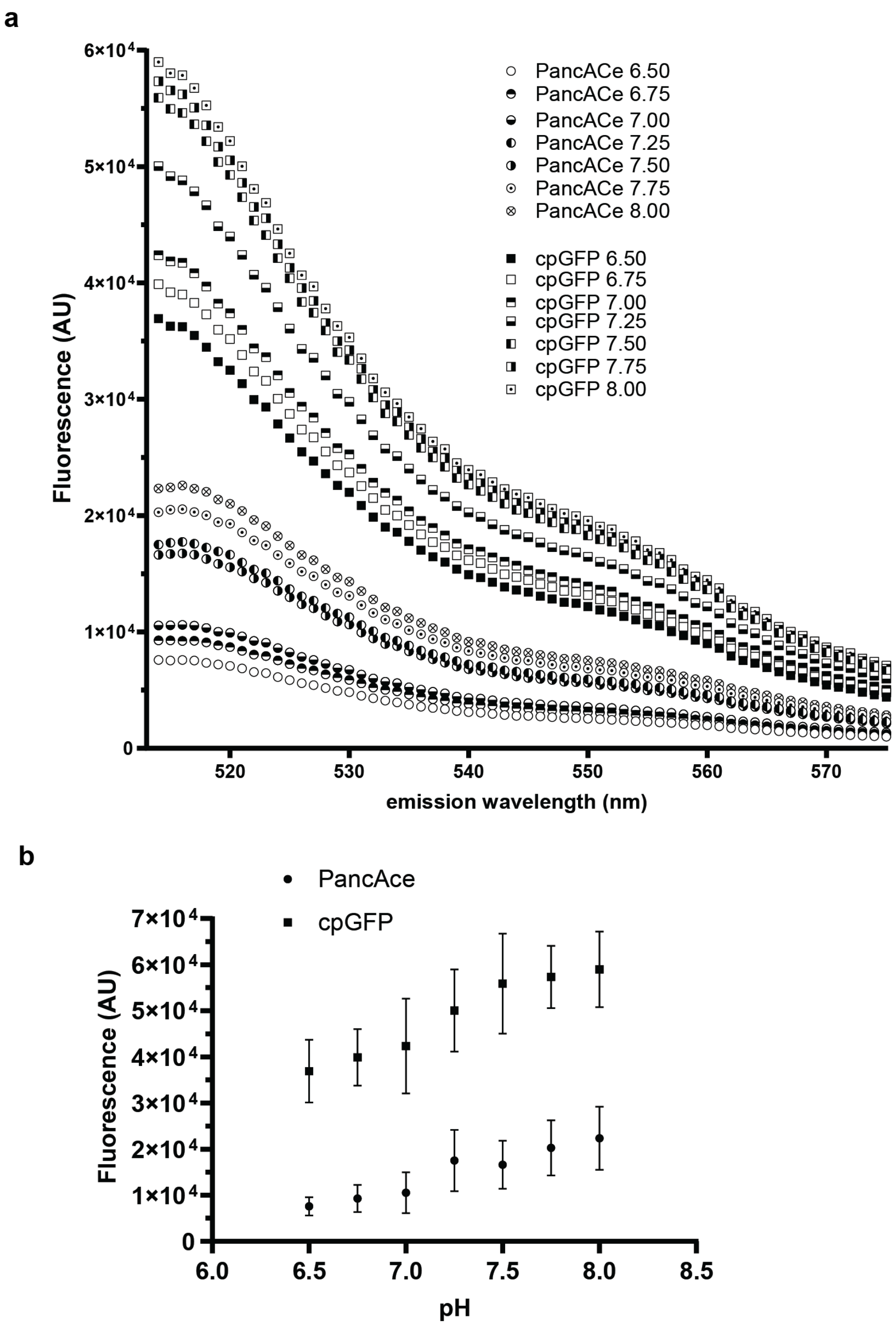


**Fig. S6. pH dependence of PancACe and cpGFP.** a) Emission spectra from pH titration of PancACe and cpGFP, $\lambda_{ex}$ = 485 nm. b) pH titration of PancACe and cpGFP at $\lambda_{em}$ = 514 nm (from panel a), n = 3, SD is shown.

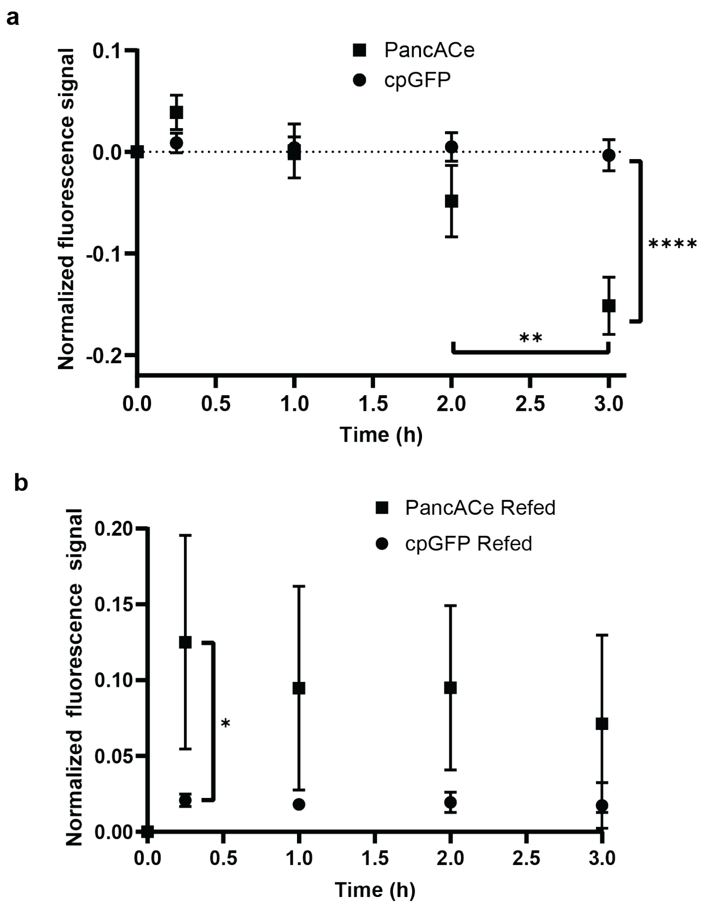


**Fig. S7. Flow cytometry analysis of PancACe and cpGFP-expressing E. coli.** a) Normalized fluorescence response of PancACe or cpGFP in *E. coli* to glucose deprivation. The cells were deprived of glucose for the indicated periods of time. b) Normalized fluorescence response of PancACe or cpGFP in *E. coli* to refeeding. The cells were deprived of glucose for 3 h prior. The cells were then given 28 mM glucose for the indicated periods of time. For both panels, the fluorescence was normalized by dividing $F_{488}/F_{405}$ and then $(F_1-F_0)/F_0$ where $F_0$ is defined by the fluorescence from cells before starvation or refeeding. $\lambda_{ex}$ = 405,485, $\lambda_{em}$ = 514 nm, n = 3, SD shown, Tukey's multiple comparison test, *p<0.05, **p<0.01, ****p<0.0001. All of the p values are listed in **Tables S4-S5.**

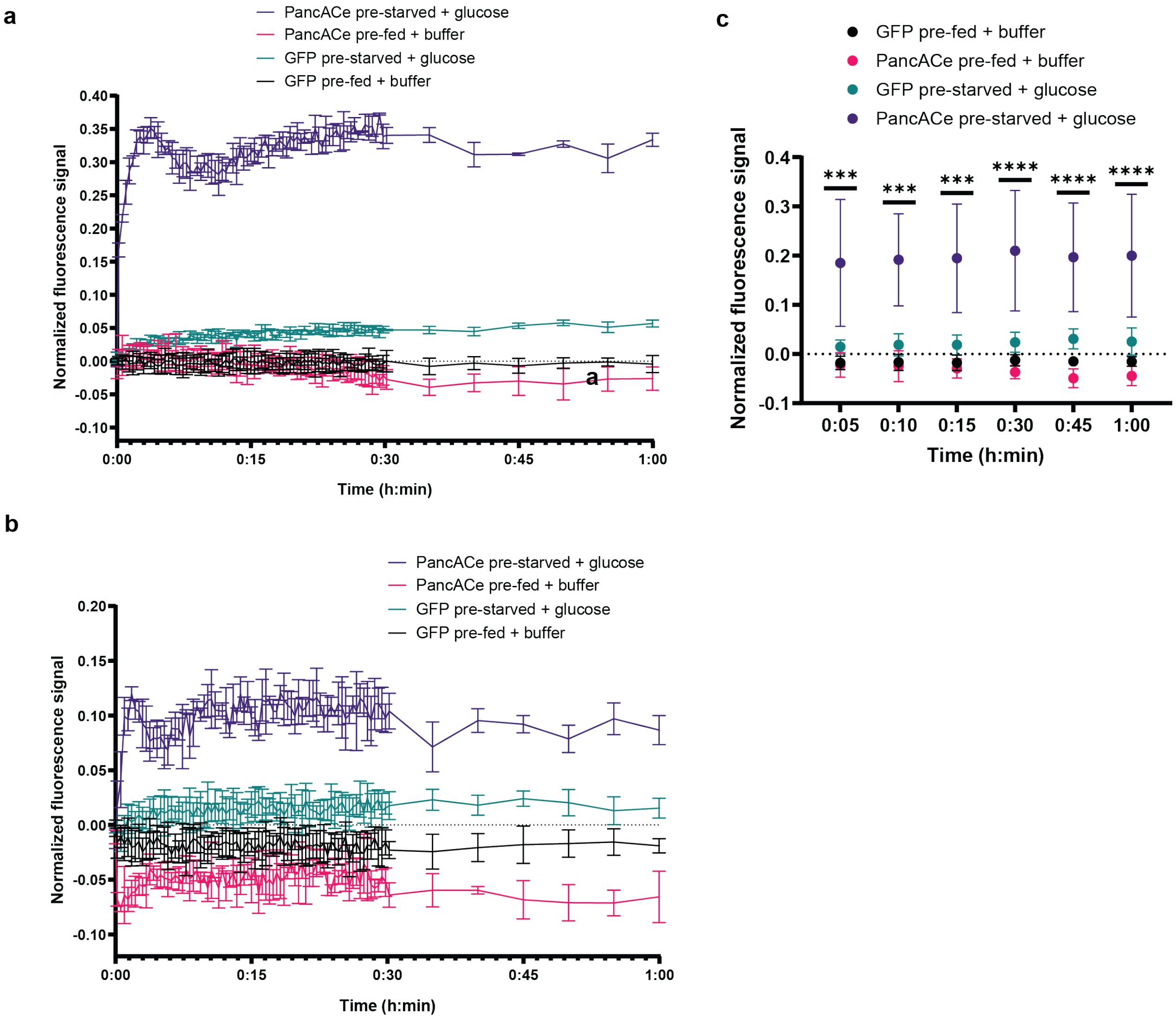


**Fig. S8**. **Plate reader analysis of glucose refeeding of PancACe and cpGFP-expressing E. coli.** a,b) Replicates of the experiment shown in **Figure 3a** performed across different days. The cells were either fed 28 mM glucose for 3 h prior ("pre-fed") or deprived of glucose for 3 h prior ("pre-starved"). At time 0:00, the cells were given either buffer alone ("+buffer") or 28 mM glucose ("+glucose"). The fluorescence was normalized by dividing $F_{488}/F_{405}$ and then $(F_1\text{-}F_0)/F_0$ where $F_0$ is defined by the fluorescence from each population of cells prior to time 0:00. $\lambda_{ex}$ = 405,485, $\lambda_{em}$ = 514 nm, n = 3 (technical replicates), SD shown. c) Averaging of the data from the selected time points across the experiments in **Figure 3a, S8a,** and **S8b**, n = 3 (experimental replicates), SD shown. Tukey's multiple comparisons test was used. The indicated p values on the plot refer to "PancACe pre-fed + buffer" compared to "PancACe pre-starved + glucose." ***p<0.001, ****p<0.0001. All of the p values are listed in **Table S6.**

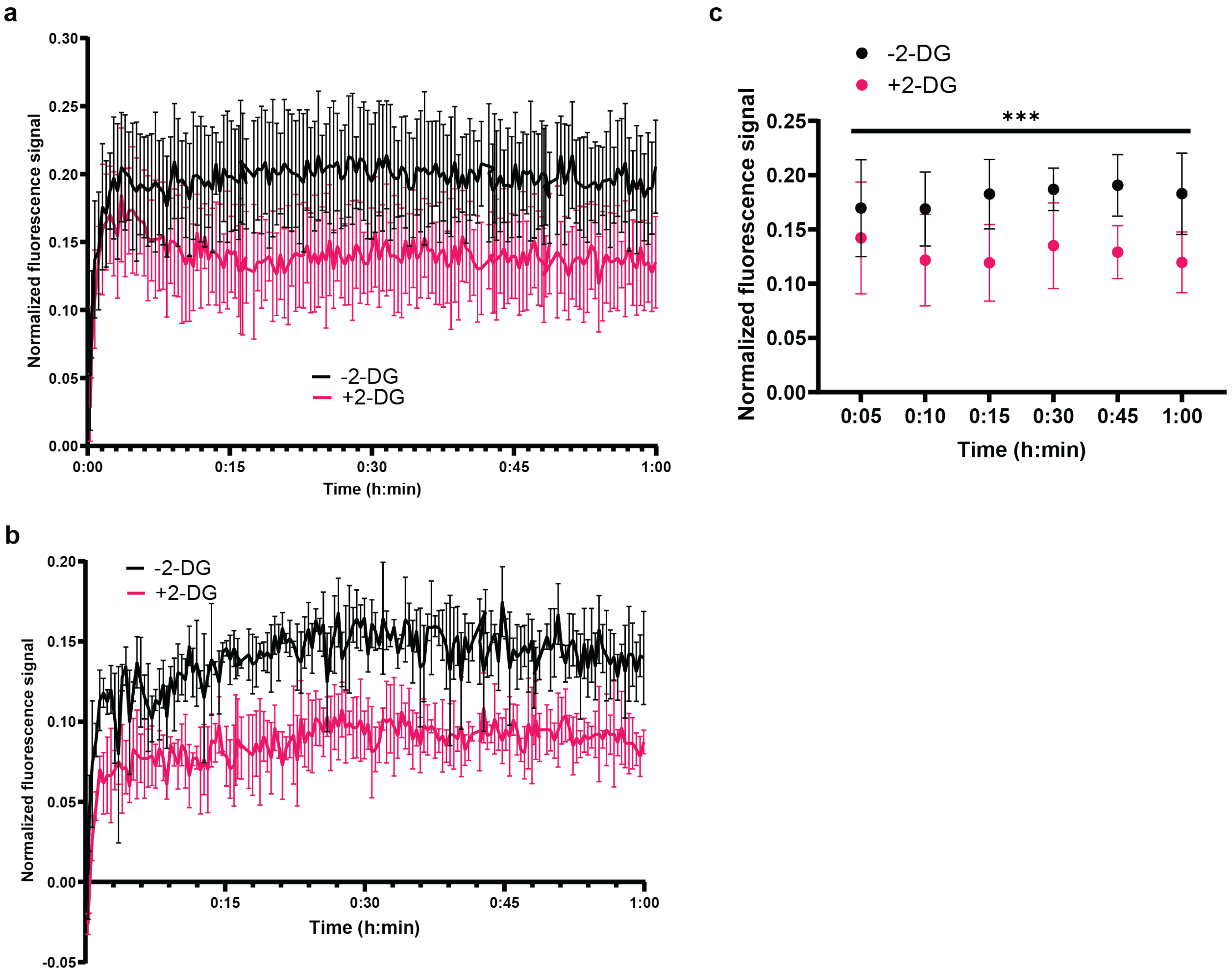


**Fig. S9. Plate reader analysis of refeeding of PancACe and cpGFP-expressing E. coli with and without 2-DG.** a) Replicates of the experiment shown in **Figure 3b** performed across different days. The cells were deprived of glucose for 3 h prior with or without 2-DG. At time 0:00, the cells were given 28 mM glucose. The fluorescence was normalized by dividing $F_{488}/F_{405}$ and then $(F_1\text{-}F_0)/F_0$ where $F_0$ is defined by the fluorescence from each population of cells prior to time 0:00. $\lambda_{ex}$ = 405,485, $\lambda_{em}$ = 514 nm, n = 3 (technical replicates), SD shown. c) Averaging of the data from the selected time points across the experiments in **Figure 3b, S9a,** and **S9b**, n = 3 (experimental replicates), SD shown, Two-way ANOVA was used, p = 0.0002. All of the p values are listed in **Table S7**.

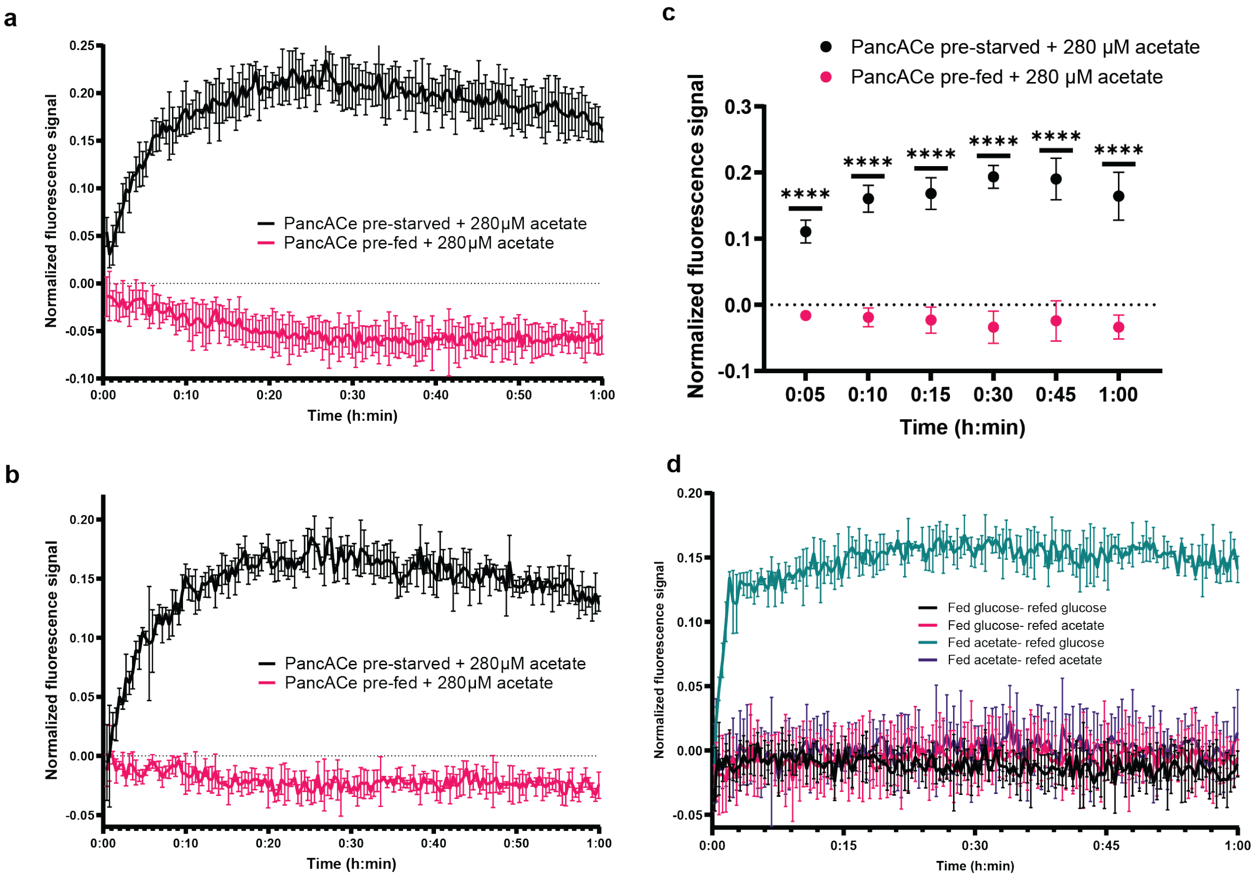


**Fig. S10. Plate reader analysis of acetate refeeding of PancACe and cpGFP-expressing E. coli.** a,b) Replicates of the experiment shown in **Figure 3c** performed across different days. The cells were either fed 28 mM glucose for 3 h prior (“pre-fed”) or deprived of glucose for 3 h prior (“pre-starved”). At time 0:00, the cells were given 280 μM acetate (“+acetate”). The fluorescence was normalized by dividing $F_{488}/F_{405}$ and then $(F_1-F_0)/F_0$ where $F_0$ is defined by the fluorescence from each population of cells prior to time 0:00. $\lambda_{ex}$ = 405,485, $\lambda_{em}$ = 514 nm, n = 3 (technical replicates), SD shown. c) Averaging of the data from the selected time points across the experiments in **Figure 3c, S10a,** and **S10b**, n = 3 (experimental replicates), SD shown, Tukey’s multiple comparisons test was used,****$p<0.0001$. All of the p values are listed in **Table S8**. d) Normalized fluorescence response of PancACe or cpGFP in *E. coli* to refeeding with glucose or acetate. The cells were either fed 28 mM glucose (“fed glucose”) or 280 μM acetate (“fed acetate) for 3 h prior. At time 0:00, the cells were given either 28 mM glucose (“+glucose”) or 280 μM acetate (“+acetate”). The fluorescence was normalized by dividing $F_{488}/F_{405}$ and then $(F_1-F_0)/F_0$ where $F_0$ is defined by the fluorescence from each population of cells prior to time 0:00. $\lambda_{ex}$ = 405,485, $\lambda_{em}$ = 514 nm, n = 3 (technical replicates), SD shown.

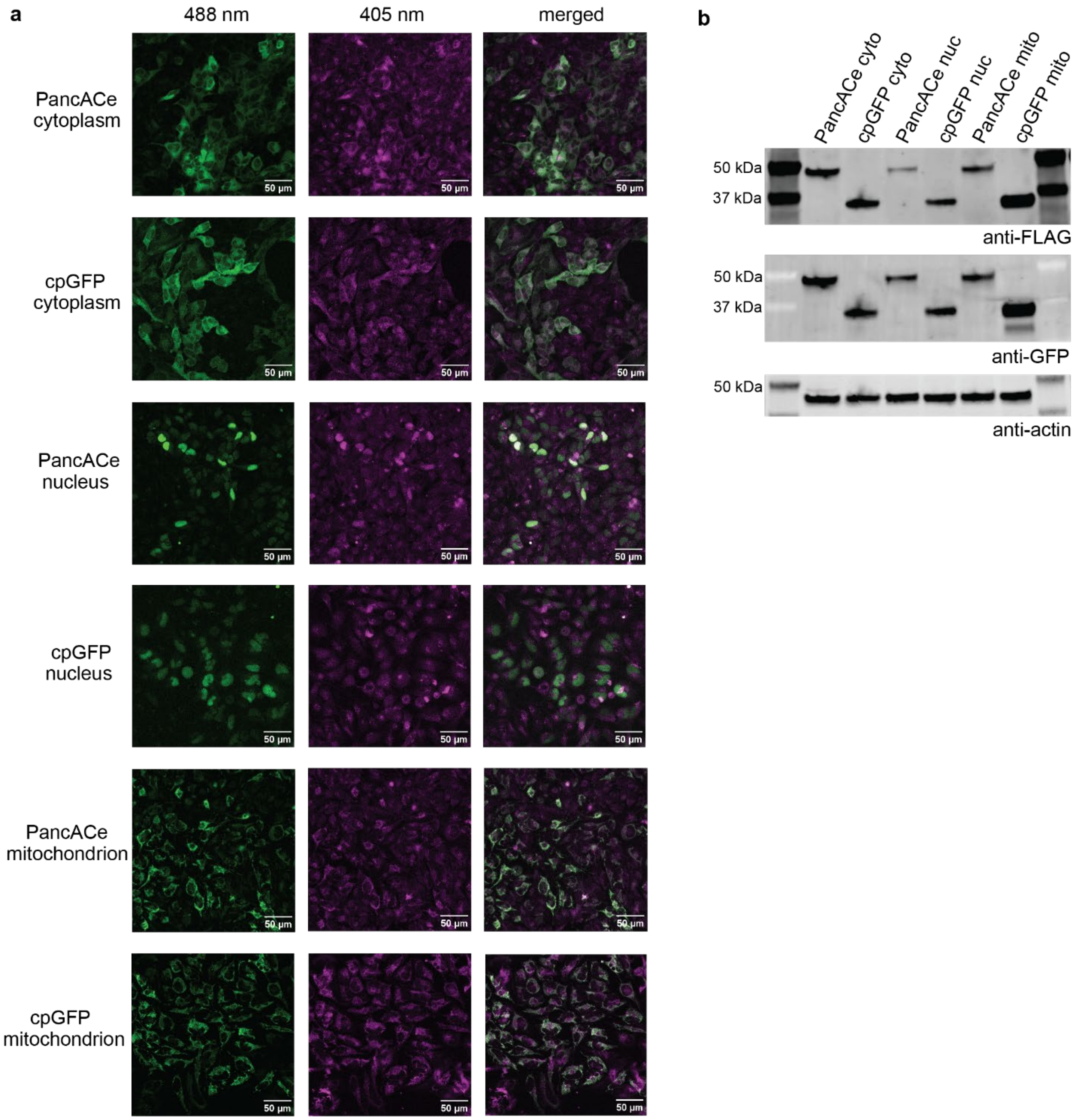


**Fig. S11. PancACe- and cpGFP-expressing HeLas.** a) Representative images for each cell line. Green is the $\lambda_{ex}$ = 488, and magenta is the $\lambda_{ex}$ = 405. b) Immunoblot analysis of lysates from the six cell lines. The lysates were analyzed on a 4-12% Bis Tris gel and probed with anti-FLAG and anti-GFP to indicate PancACe or cpGFP. Anti-actin was used as the loading control.

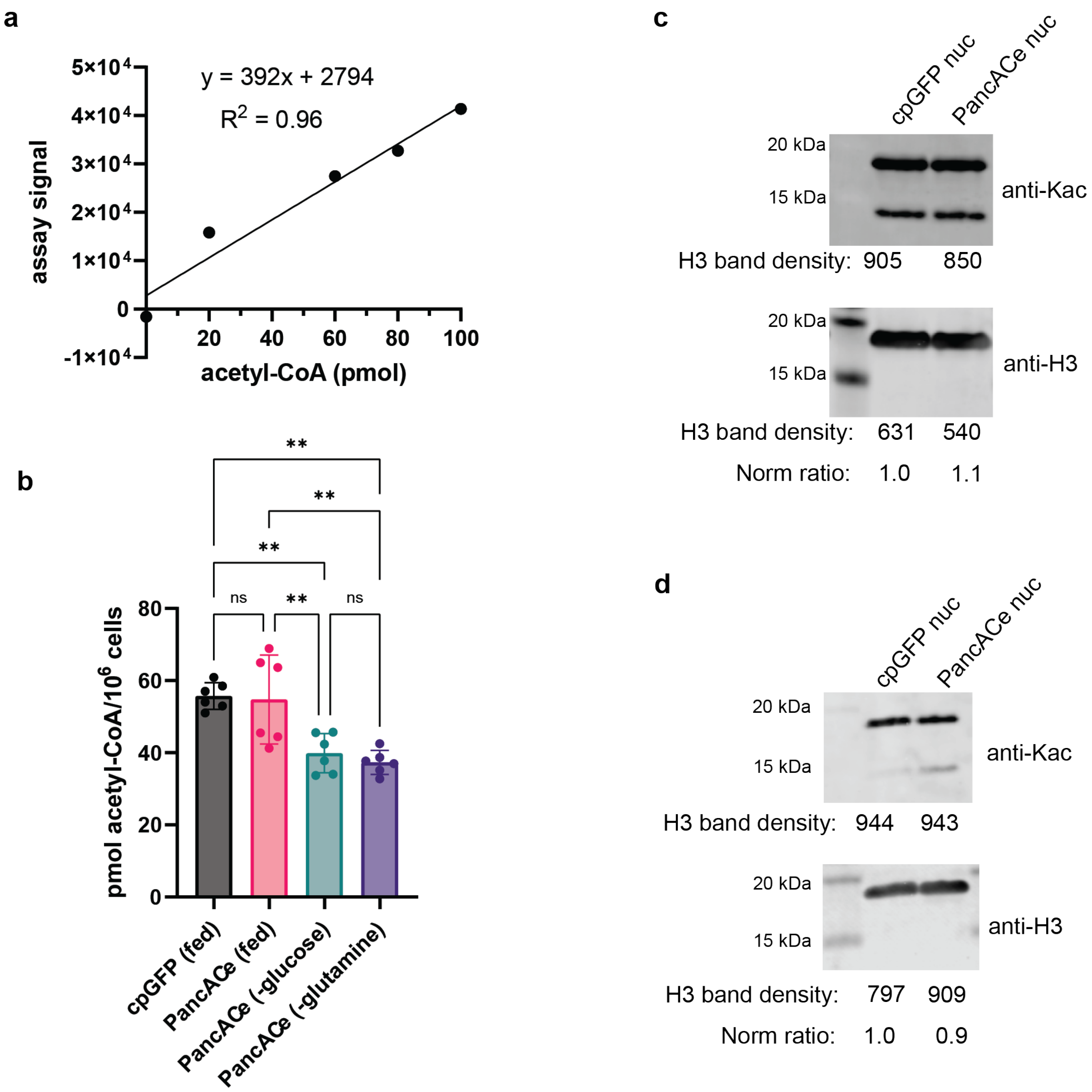


**Fig. S12. PicoProbe™ analysis of whole cell free acetyl-CoA and immunoblot analysis of histone acetylation.** a) Calibration curve generated using the PicoProbe™ kit (n = 1). The data were fitted to a line with the equation shown in the inset. b) Acetyl-CoA in the cell lysate samples from the cyto-cpGFP or cyto-PancACe HeLa cell lines as determined by PicoProbe™ kit (n = 6). The "-glucose" and "-glutamine" cells were deprived of these nutrients for 16 h before lysis. SD shown, Tukey's multiple comparison test was used, **p<0.01. All of the p values are listed in **Table S9.** c,d) Immunoblot analysis of lysine acetylation (anti-Kac) of extracted histones from nuc-cpGFP and nuc-PancACe cells (n = 2). The samples were analyzed using a 12% Bis Tris gel. H3 WT was used as the loading control. The densitometry values from Licor Image Studio are shown.

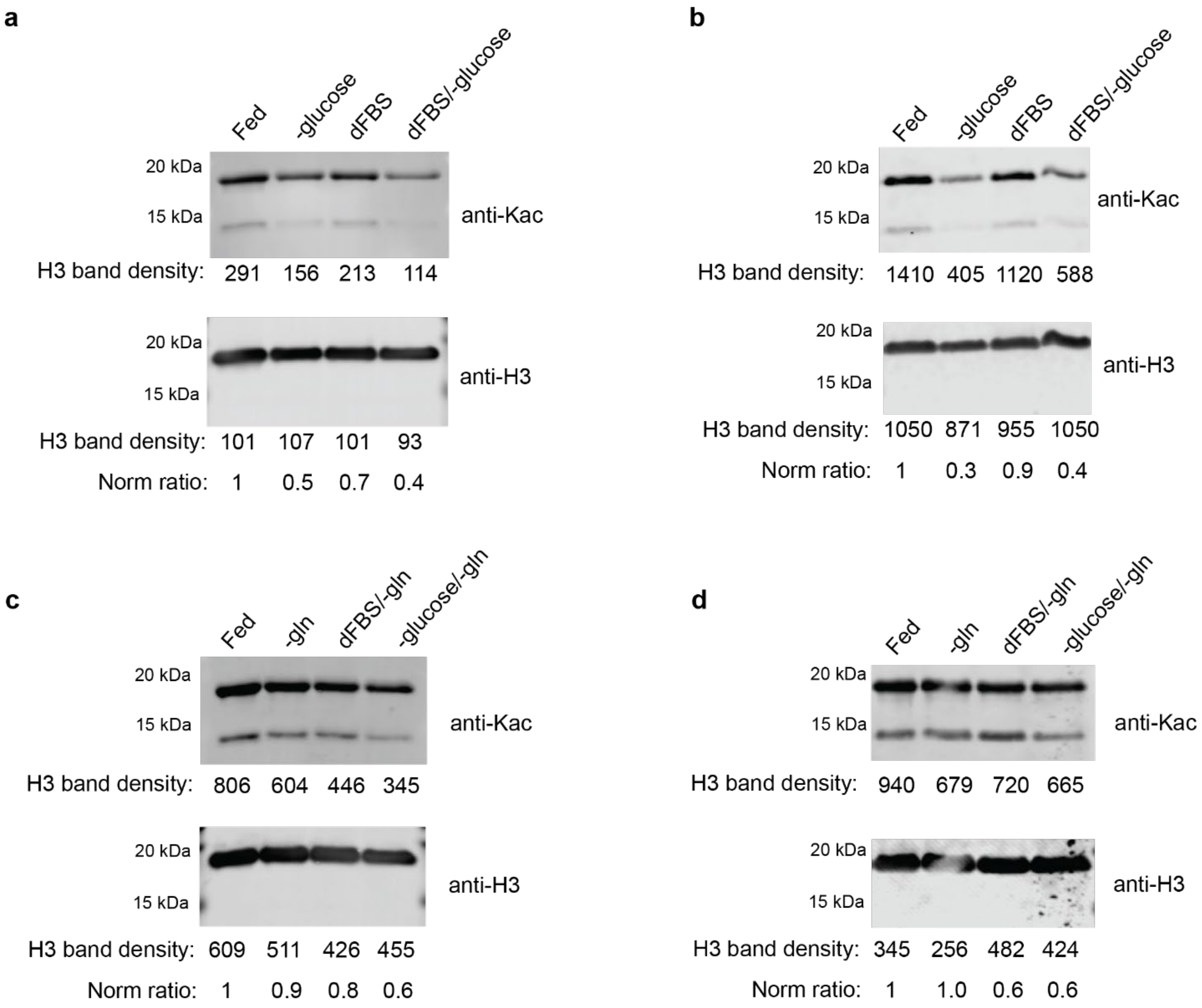


**Fig. S13. Immunoblot analysis of lysine acetylation of extracted histones from deprived HeLas.** a-d) Anti-Kac blots of extracted histones from nuc-cpGFP cells that were subjected to 16 h of the indicated deprivation conditions (n = 2). H3 WT was used as the loading control. The densitometry values from Licor Image Studio are shown.

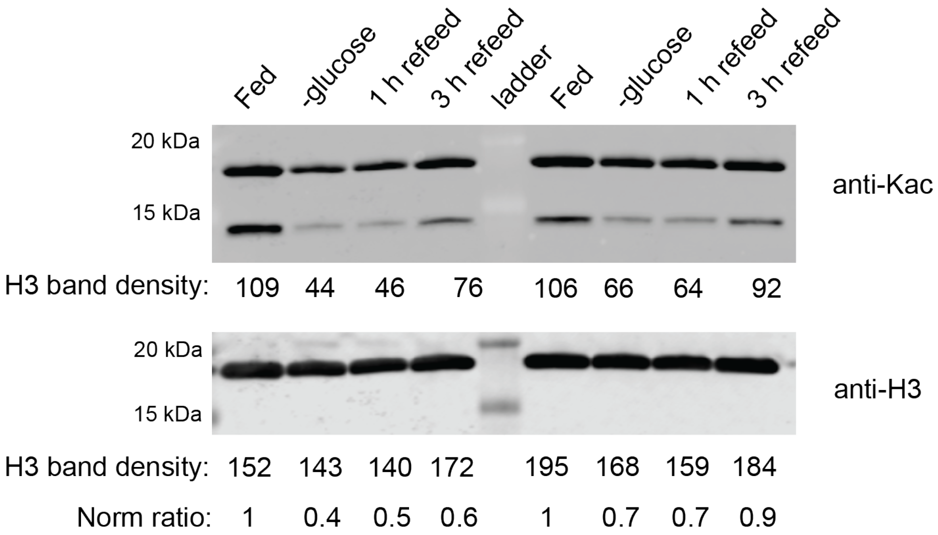


**Fig. S14. Immunoblot analysis of refed HeLas.** Anti-Kac blots of extracted histones from nuc-cpGFP cells that were subjected to 16 h of glucose deprivation and then refed with glucose for 1 h or 3 h (n = 2). H3 WT was used as the loading control. The densitometry values from Licor Image Studio are shown.

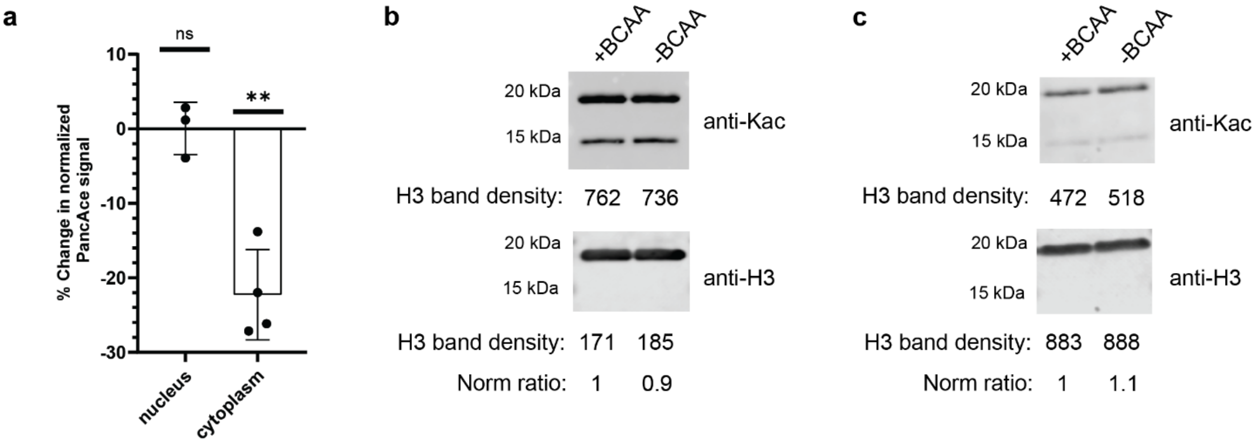


**Fig. S15. Analysis of +/-BCAA-treated HeLas.** a) PancACe signal derived from images of HeLa cells expressing nuc-PancACe or cyto-PancACe that were deprived of BCAAs for 24 h normalized to cells that received BCAAs during that time period. The data were collected and normalized as described in the Methods section. n = 3-4 experimental replicates, SD shown. Two-tailed, paired t tests were performed to compare -/+ BCAA cells: nuc $p = 0.8898$, cyto $p = 0.0091$, b-c) Anti-Kac blots of extracted histones from nuc-cpGFP cells that were given +/-BCAA for 24 h (n = 2). H3 WT was used as the loading control. The densitometry values from Licor Image Studio are shown.

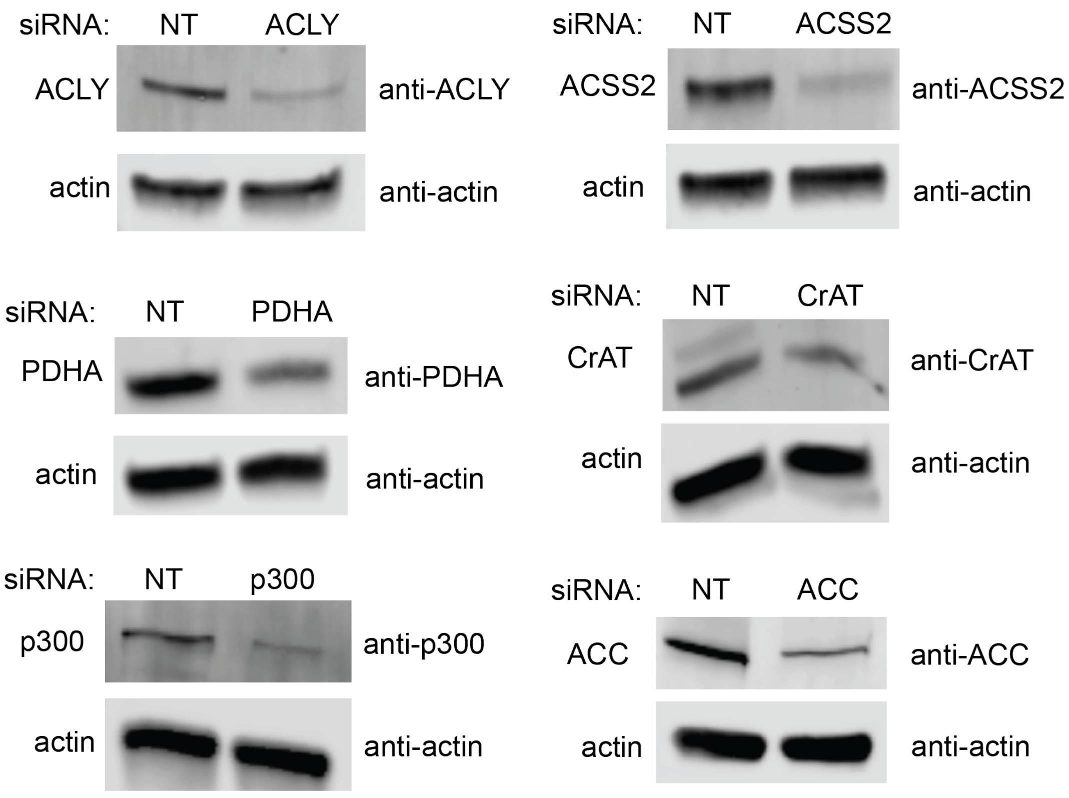

**Fig. S16. Immunoblot validation of siRNA knockdowns in HeLas.** Lysates from HeLa cell lines transfected with non-targeted (NT) or targeted siRNAs for 48 h were analyzed with protein specific antibodies to assess the knockdown of each target as shown (n = 1). Anti-actin was used as a loading control.

a. Figure S11b

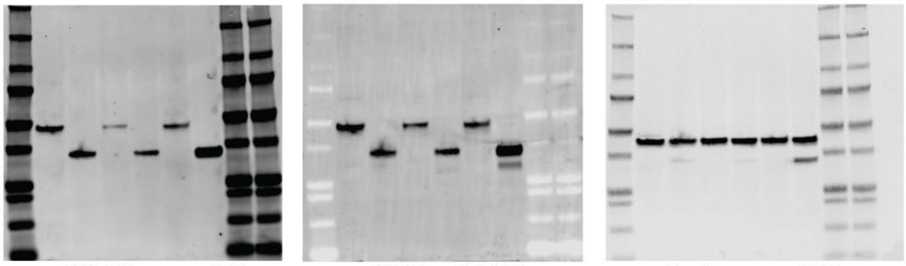

anti-FLAG, mouse-680 anti-GFP, rabbit-800 anti-actin, mouse-680

b. Figure S12d Figure S15c Figure S13b Figure S12d Figure S15c Figure S13b

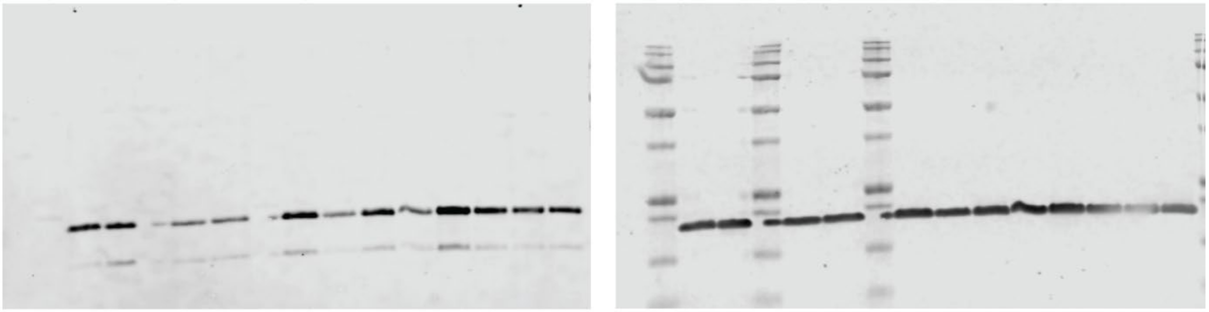

anti-Kac, rabbit-800 anti-H3, mouse-680

c. Figure S12c

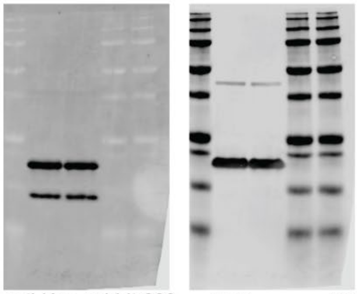

anti-Kac, rabbit-800 anti-H3, mouse-680

d. Figure S13a

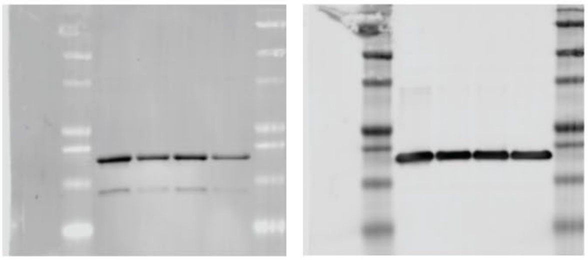

anti-Kac, rabbit-800 anti-H3, mouse-680

e. Figure S13c

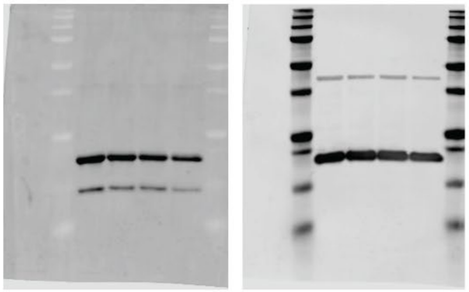

anti-Kac, rabbit-800 anti-H3, mouse-680

f. Figure S13d

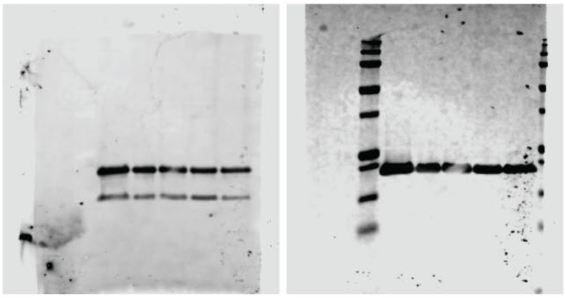

anti-Kac, rabbit-800 anti-H3, mouse-680

g. Figure S14

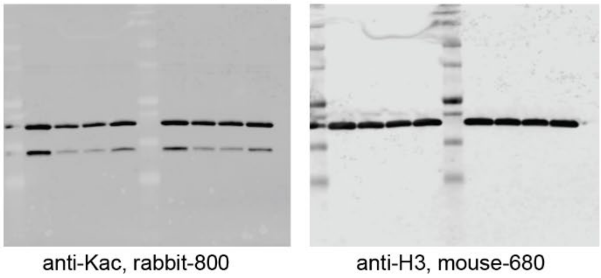


h. Figure S15b

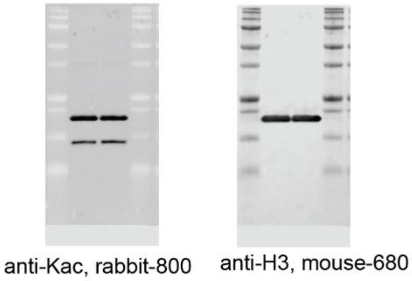


i. Figure S16

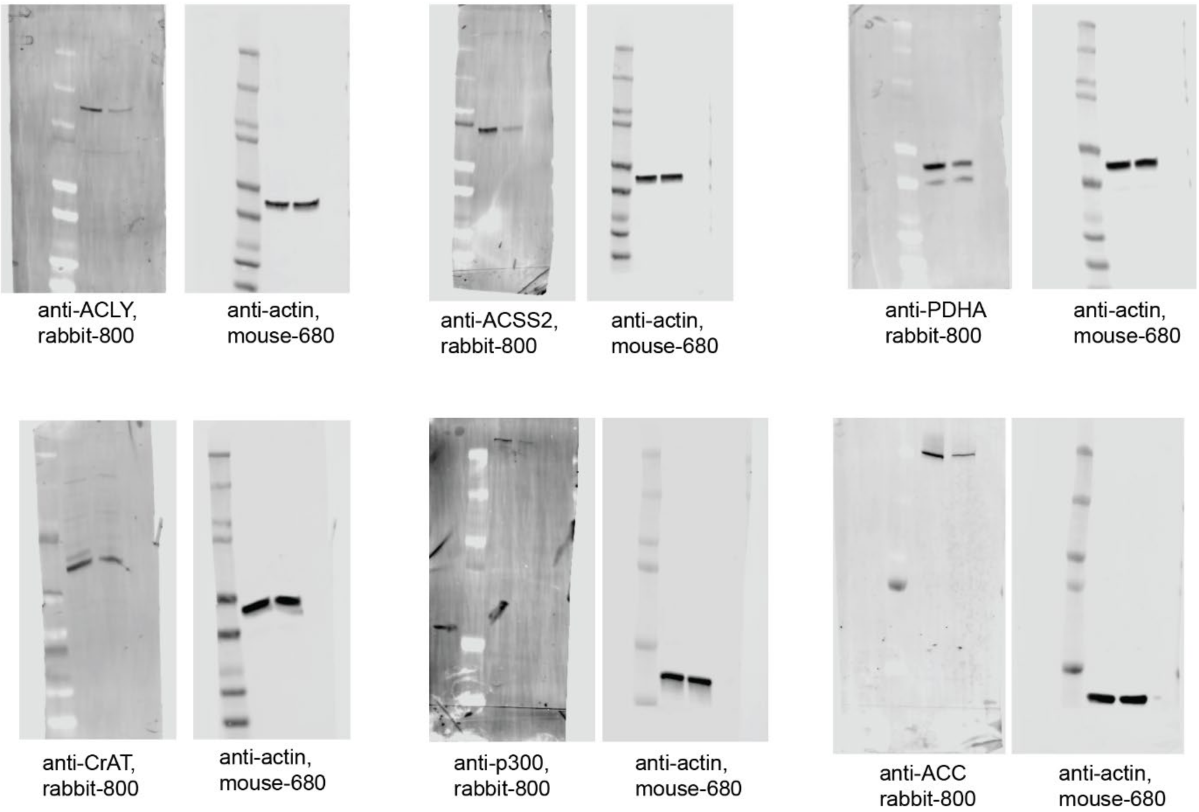


**Fig. S17**. **Uncropped blots.** The entire lane is shown in each image. a) Blots in S11b, b) Blots in S12d, S15c, and S13b. Note that the right side of the blot was not used due to the transfer artifact that is apparent in the mouse-680 for that sample set. c) Blots in S12c, d) Blots in S13a, e) Blots in S13c, f) Blots in S13d. Note that the first lane to the right of the ladder was accidentally loaded with two different samples and so was disregarded. g) Blots in S14, h) Blots in S15b, i) Blots in S16.

**Table S1.** *p values from two tailed, paired t test of HeLa cell deprivation experiments*

| **Condition comparison** | **Cytoplasm** | **Nucleus** | **Mitochondrion** |
|---|---|---|---|
| Fed vs -glucose | *0.0181* | *0.0087* | 0.9185 |
| Fed vs dFBS | *0.0411* | 0.4145 | 0.2103 |
| Fed vs dFBS/-glucose | *0.0054* | *0.0171* | 0.1011 |
| Fed vs -glutamine | *0.0012* | *0.0227* | 0.3526 |
| Fed vs dFBS/-glutamine | *0.0184* | 0.1948 | 0.8184 |
| Fed vs -glucose/-glutamine | *<0.0001* | *0.0016* | *0.0330* |

**Table S2.** *p values from HeLa cell refeeding experiments.* Comparison to Fed are two-tailed, paired t tests, while comparison between other conditions are from Tukey's multiple comparison test

| **Condition comparison** | **Cytoplasm** | **Nucleus** |
|---|---|---|
| Fed vs -gluc | *0.0246* | *0.0082* |
| Fed vs +gluc 1 h | 0.1185 | 0.2026 |
| Fed vs +gluc 3 h | 0.2534 | 0.1227 |
| -Gluc vs. +gluc 1 h | 0.8791 | *0.0037* |
| -Gluc vs. +gluc 3 h | *0.0298* | *0.0152* |
| +Gluc 1 h vs. +gluc 3 h | 0.0542 | 0.7528 |

**Table S3.** *p values from two tailed, paired t test of HeLa cell siRNA experiments*

| Condition comparison | Cytoplasm | Nucleus | Mitochondrion |
|---|---|---|---|
| NT vs ACLY | 0.0473 | 0.117 | 0.3195 |
| NT vs ACSS2 | 0.0012 | 0.0016 | 0.4224 |
| NT vs PDHA | 0.3002 | 0.0202 | 0.3937 |
| NT vs CrAT | 0.0677 | 0.0512 | 0.6914 |
| NT vs p300 | 0.0694 | 0.0864 | 0.1646 |
| NT vs ACC | 0.0193 | 0.0879 | 0.0194 |

**Table S4.** *p values from Tukey's multiple comparison of the E. coli glucose starvation FACS experiment*

| Condition Comparison | p value |
|---|---|
| GFP 0.25 min vs. Pan 0.25 min | 0.7034 |
| GFP 0.25 min vs. GFP 1 h | >0.9999 |
| GFP 0.25 min vs. Pan 1 h | 0.9998 |
| GFP 0.25 min vs. GFP 2 h | >0.9999 |
| GFP 0.25 min vs. Pan 2h | 0.0902 |
| GFP 0.25 min vs. GFP 3 h | 0.9964 |
| GFP 0.25 min vs. Pan 3 h | <0.0001 |
| Pan 0.25 min vs. GFP 1 h | 0.5443 |
| Pan 0.25 min vs. Pan 1 h | 0.4528 |
| Pan 0.25 min vs. GFP 2 h | 0.5739 |
| Pan 0.25 min vs. Pan 2h | 0.0046 |
| Pan 0.25 min vs. GFP 3 h | 0.3340 |
| Pan 0.25 min vs. Pan 3 h | <0.0001 |
| GFP 1 h vs. Pan 1 h | >0.9999 |
| GFP 1 h vs. GFP 2 h | >0.9999 |
| GFP 1 h vs. Pan 2h | 0.1427 |
| GFP 1 h vs. GFP 3 h | 0.9999 |
| GFP 1 h vs. Pan 3 h | <0.0001 |
| Pan 1 h vs. GFP 2 h | >0.9999 |
| Pan 1 h vs. Pan 2h | 0.1853 |
| Pan 1 h vs. GFP 3 h | >0.9999 |
| Pan 1 h vs. Pan 3 h | <0.0001 |
| GFP 2 h vs. Pan 2h | 0.1313 |
| GFP 2 h vs. GFP 3 h | 0.9997 |
| GFP 2 h vs. Pan 3 h | <0.0001 |
| Pan 2h vs. GFP 3 h | 0.2656 |
| Pan 2h vs. Pan 3 h | 0.0010 |
| GFP 3 h vs. Pan 3 h | <0.0001 |

**Table S5.** *p values from Tukey's multiple comparison of the E. coli glucose refeeding FACS experiment*

| **Condition comparison** | **p value** |
|---|---|
| GFP 0.25 vs. GFP 1 | >0.9999 |
| GFP 0.25 vs. GFP 2 h | >0.9999 |
| GFP 0.25 vs. GFP 3 h | >0.9999 |
| GFP 0.25 vs. Pan 0.25 | 0.0183 |
| GFP 0.25 vs. Pan 1 | 0.1456 |
| GFP 0.25 vs. Pan 2 | 0.1435 |
| GFP 0.25 vs. Pan 3 h | 0.5256 |
| GFP 1 vs. GFP 2 h | >0.9999 |
| GFP 1 vs. GFP 3 h | >0.9999 |
| GFP 1 vs. Pan 0.25 | 0.0151 |
| GFP 1 vs. Pan 1 | 0.1227 |
| GFP 1 vs. Pan 2 | 0.1210 |
| GFP 1 vs. Pan 3 h | 0.4674 |
| GFP 2 h vs. GFP 3 h | >0.9999 |
| GFP 2 h vs. Pan 0.25 | 0.0166 |
| GFP 2 h vs. Pan 1 | 0.1334 |
| GFP 2 h vs. Pan 2 | 0.1316 |
| GFP 2 h vs. Pan 3 h | 0.4955 |
| GFP 3 h vs. Pan 0.25 | 0.0144 |
| GFP 3 h vs. Pan 1 | 0.1172 |
| GFP 3 h vs. Pan 2 | 0.1155 |
| GFP 3 h vs. Pan 3 h | 0.4521 |
| Pan 0.25 vs. Pan 1 | 0.9222 |
| Pan 0.25 vs. Pan 2 | 0.9248 |
| Pan 0.25 vs. Pan 3 h | 0.4523 |
| Pan 1 vs. Pan 2 | >0.9999 |
| Pan 1 vs. Pan 3 h | 0.9785 |
| Pan 2 vs. Pan 3 h | 0.9774 |

**Table S6.** *p values from Tukey's multiple comparison of the E. coli glucose refeeding experiment*

| **Condition Comparison** | **p value** |
|---|---|
| 0:05 | 0.9998 |
| GFP pre-fed + buffer vs. PancACe pre-fed + buffer | 0.9098 |
| GFP pre-fed + buffer vs. GFP pre-starved + glucose | 0.0008 |
| GFP pre-fed + buffer vs. PancACe pre-starved + glucose | 0.8760 |
| PancACe pre-fed + buffer vs. GFP pre-starved + glucose | 0.0006 |
| PancACe pre-fed + buffer vs. PancACe pre-starved + glucose | 0.0059 |
| GFP pre-starved + glucose vs. PancACe pre-starved + glucose | |
| 0:10 | 0.9986 |
| GFP pre-fed + buffer vs. PancACe pre-fed + buffer | 0.8873 |
| GFP pre-fed + buffer vs. GFP pre-starved + glucose | 0.0006 |
| GFP pre-fed + buffer vs. PancACe pre-starved + glucose | 0.8144 |
| PancACe pre-fed + buffer vs. GFP pre-starved + glucose | 0.0003 |
| PancACe pre-fed + buffer vs. PancACe pre-starved + glucose | 0.0052 |
| GFP pre-starved + glucose vs. PancACe pre-starved + glucose | |
| 0:15 | 0.9956 |
| GFP pre-fed + buffer vs. PancACe pre-fed + buffer | 0.8789 |
| GFP pre-fed + buffer vs. GFP pre-starved + glucose | 0.0004 |
| GFP pre-fed + buffer vs. PancACe pre-starved + glucose | 0.7646 |
| PancACe pre-fed + buffer vs. GFP pre-starved + glucose | 0.0002 |
| PancACe pre-fed + buffer vs. PancACe pre-starved + glucose | 0.0043 |
| GFP pre-starved + glucose vs. PancACe pre-starved + glucose | |
| 0:30 | 0.9608 |
| GFP pre-fed + buffer vs. PancACe pre-fed + buffer | 0.8779 |
| GFP pre-fed + buffer vs. GFP pre-starved + glucose | 0.0002 |
| GFP pre-fed + buffer vs. PancACe pre-starved + glucose | 0.6071 |
| PancACe pre-fed + buffer vs. GFP pre-starved + glucose | <0.0001 |
| PancACe pre-fed + buffer vs. PancACe pre-starved + glucose | 0.0023 |
| GFP pre-starved + glucose vs. PancACe pre-starved + glucose | |
| 0:45 | 0.8980 |
| GFP pre-fed + buffer vs. PancACe pre-fed + buffer | 0.7894 |
| GFP pre-fed + buffer vs. GFP pre-starved + glucose | 0.0005 |
| GFP pre-fed + buffer vs. PancACe pre-starved + glucose | 0.3744 |
| PancACe pre-fed + buffer vs. GFP pre-starved + glucose | <0.0001 |
| PancACe pre-fed + buffer vs. PancACe pre-starved + glucose | 0.0077 |
| GFP pre-starved + glucose vs. PancACe pre-starved + glucose | |
| 1:00 | 0.9325 |
| GFP pre-fed + buffer vs. PancACe pre-fed + buffer | 0.8412 |
| GFP pre-fed + buffer vs. GFP pre-starved + glucose | 0.0004 |
| GFP pre-fed + buffer vs. PancACe pre-starved + glucose | 0.4908 |
| PancACe pre-fed + buffer vs. GFP pre-starved + glucose | <0.0001 |
| PancACe pre-fed + buffer vs. PancACe pre-starved + glucose | 0.0047 |
| GFP pre-starved + glucose vs. PancACe pre-starved + glucose | 0.9998 |

**Table S7.** *p values from Tukey's multiple comparison of the E. coli +/-2-DG glucose refeeding experiment*

| **Condition comparison** | **p value** |
|---|---|
| 0:05 | |

| | |
|---|---|
| -2-DG vs. +2-DG | 0.3573 |
| 0:10 | |
| -2-DG vs. +2-DG | 0.1191 |
| 0:15 | |
| -2-DG vs. +2-DG | 0.0404 |
| 0:30 | |
| -2-DG vs. +2-DG | 0.0883 |
| 0:45 | |
| -2-DG vs. +2-DG | 0.0457 |
| 1:00 | |
| -2-DG vs. +2-DG | 0.0406 |

**Table S8.** *p values from Tukey's multiple comparison of the E. coli acetate refeeding experiment*

| Condition Comparison | p value |
|---|---|
| 0:05 | |
| PancACe pre-starved + acetate vs. PancACe pre-fed + acetate | <0.0001 |
| 0:10 | |
| PancACe pre-starved + acetate vs. PancACe pre-fed + acetate | <0.0001 |
| 0:15 | |
| PancACe pre-starved + acetate vs. PancACe pre-fed + acetate | <0.0001 |
| 0:30 | |
| PancACe pre-starved + acetate vs. PancACe pre-fed + acetate | <0.0001 |
| 0:45 | |
| PancACe pre-starved + acetate vs. PancACe pre-fed + acetate | <0.0001 |
| 1:00 | |
| PancACe pre-starved + acetate vs. PancACe pre-fed + acetate | <0.0001 |

**Table S9.** *p values from Tukey's multiple comparison of the PicoProbe data*

| Condition comparison | p value |
|---|---|
| cpGFP (fed) vs. PancACe (fed) | 0.9955 |
| cpGFP (fed) vs. PancACe (-glucose) | 0.0053 |
| cpGFP (fed) vs. PancACe (-glutamine) | 0.0013 |
| PancACe (fed) vs. PancACe (-glucose) | 0.0089 |
| PancACe (fed) vs. PancACe (-glutamine) | 0.0022 |
| PancACe (-glucose) vs. PancACe (-glutamine) | 0.9251 |

**Table S10.** *Results of the energy minimization of nineteen linker composition variants.* The top model has been highlighted in red (GE).

| Amino Acids | YASARA Energy (kJ/mol) | Amino Acids | YASARA Energy (kJ/ mol) | Amino Acids | YASARA Energy (kJ/ mol) | Amino Acids | YASARA Energy (kJ/ mol) |
|---|---|---|---|---|---|---|---|
| GG | -195702.3 | GV | -196706.3 | GH | -196730.5 | ***GE*** | ***-197975.8*** |
| GL | -195967.1 | GS | -195952.5 | GW | -196204.4 | GN | -195617.2 |
| GI | -196323.0 | GT | -196499.4 | GK | -195986.9 | GQ | -197225.4 |
| GM | -196044.0 | GP | -195838.7 | GR | -195927.6 | GC | -195132.2 |
| GF | -195564.1 | GY | -197197.7 | GD | -196233.4 | **GA (ref.)** | **-196993.7** |

**Table S11**: *Results of the energy minimization of specific repeat linker sequences and reversed sequences.* The top model has been highlighted in red (EE).

| Amino Acid | YASARA Energy (kJ/mol) |
|---|---|
| AA | -195005.3 |
| AG | -195790.0 |
| ***EE*** | ***-199736.3*** |
| EG | -198319.1 |
| QG | -196535.0 |
| QQ | -197844.1 |

**Table S12**. *Docking results of the linker variants, compared to PancACe (XY = GA).*

| **Linker** | **Docking Score (kcal/mol)** |
|---|---|
| GA (Original) | -14.20 |
| GG (control) | -12.46 |
| GE | -13.64 |
| EG | -12.64 |
| QQ | -14.03 |
| GQ | -14.40 |
| EE | -14.56 |

**Table S13.** *Media formulations for deprivation experiments in HeLa cells*

| Condition | Glucose (A24940-01, Gibco) | Glutamine (Gibco, 25030081) | FBS or dFBS (dFBS: A33820-01, Gibco) |
|---|---|---|---|
| Fed | 4.5 g/L | 4 mM | 10% v/v FBS |
| -glucose | 0 g/L | 4 mM | 10% v/v FBS |
| dFBS | 4.5 g/L | 4 mM | 10% v/v dFBS |
| dFBS/-glucose | 0 g/L | 4 mM | 10% v/v dFBS |
| -glutamine | 4.5 g/L | 0 mM | 10% v/v FBS |
| dFBS/-glutamine | 4.5 g/L | 0 mM | 10% v/v dFBS |
| -glucose/-glutamine | 0 g/L | 0 mM | 10% v/v FBS |

**Table S14.** *siRNAs*

| siRNA | Catalog ID |
|---|---|
| ON-TARGETplus Non-targeting Pool | D-001810-10 |
| ON-TARGETplus Human ACLY siRNA | L-004915-00-0005 |
| ON-TARGETplus Human PDHA1 siRNA | L-010329-00-0005 |
| ON-TARGETplus Human ACSS2 siRNA | L-010396-00-0005 |
| ON-TARGETplus Human CRAT siRNA | L-009524-00-0005 |
| ON-TARGETplus Human EP300 siRNA | L-003486-00-0005 |
| ON-TARGETplus Human ACACA siRNA | L-004551-00-0005 |
| ON-TARGETplus Human ACACB siRNA | L-004759-00-0005 |

**Table S15.** *Antibodies*

| Antibody | Manufacturer | Catalog ID |
|---|---|---|
| Rabbit anti-acetyllysine | Cell Signaling Technologies | 9441 |
| Mouse anti-histone H3 | Cell Signaling Technologies | 3638 |
| Mouse anti-actin mAb | Cell Signaling Technologies | 3700 |
| Rabbit anti-ACLY mAb | Cell Signaling Technologies | 13390 |
| Rabbit anti-ACSS2 mAb | Cell Signaling Technologies | 3658 |
| Rabbit anti-PDH mAb | Cell Signaling Technologies | 3205 |
| Rabbit anti-CrAT pAb | Proteintech | 15170-1-AP |
| Rabbit anti-p300 mAb | Cell Signaling Technologies | 57625 |
| Rabbit anti-ACC mAb | Cell Signaling Technologies | 3676 |
| IRDye® 800CW Goat anti-Rabbit IgG Secondary Antibody | Licor | 926-32211 |
| IRDye® 680RD Goat anti-Mouse IgG Secondary Antibody | Licor | 926-68070 |

**Data File S1. Fluorescence data for Figs. 1b, 1c, and S2.** This Excel file contains all of the raw fluorescence data that constitute the indicated figures. It includes a summary of the processed data as they were entered into Prism to generate the plots shown in the figures.

**Data file S2. Fluorescence data for Fig. 2b-d.** This Excel file contains all of the raw fluorescence data that constitute the indicated figures. It includes a summary of the processed data as they were entered into Prism to generate the plots shown in the figures.

**Data file S3. Fluorescence data for Fig. S6.** This Excel file contains all of the raw fluorescence data that constitute the indicated figures. It includes a summary of the processed data as they were entered into Prism to generate the plots shown in the figures.

**Data file S4. Fluorescence data for Fig 3a and S8.** This Excel files contains all of the raw and normalized fluorescence data that constitute the indicated figures.

**Data file S5. Fluorescence data for Fig 3b and S9.** This Excel files contains all of the raw and normalized fluorescence data that constitute the indicated figures.

**Data file S6. Fluorescence data for Fig 3c and S10.** This Excel files contains all of the raw and normalized fluorescence data that constitute the indicated figures.

**Data file S7. Fluorescence microscopy data from the well level.** This Excel files contains all of the data exported from SQL for the fluorescence value for each well from the microscopy experiments. The data are organized by the day each experiment was conducted. The file includes a summary of the averaged data (across all days) as they were entered into Prism to generate the plots in Figs. 4, 5, and S15. The grayed-out values were deemed to be outliers according to the Grubbs' test.
